# Intermittent Fasting Activates AMP-Kinase to Restructure Right Ventricular Lipid Metabolism and Microtubules in Two Rodent Models of Pulmonary Arterial Hypertension

**DOI:** 10.1101/2022.03.07.483333

**Authors:** Felipe Kazmirczak, Lynn M. Hartweck, Neal T. Vogel, Jenna B. Mendelson, Anna K. Park, Rashmi M. Raveendran, Jin O-Uchi, Bong Sook Jhun, Sasha Z. Prisco, Kurt W. Prins

## Abstract

Intermittent fasting (IF) extends lifespan via pleotropic mechanisms, but one important molecular mediator of the beneficial effects of IF is AMP-kinase (AMPK). AMPK enhances lipid metabolism and modulates microtubule dynamics. Dysregulation of these two molecular pathways causes right ventricular (RV) failure in pulmonary arterial hypertension (PAH). In two models of rodent PAH, we show IF activates RV AMPK, which restores mitochondrial morphology and peroxisomal density and restructures mitochondrial/peroxisomal lipid metabolism protein regulation. IF also increases electron transport chain (ETC) protein abundance and activity in the RV. Echocardiographic and hemodynamic measures of RV function are positively associated with fatty acid oxidation and ETC protein levels in correlational heatmapping analyses. IF also combats heightened microtubule density, which normalizes t-tubule structure. In summation, we demonstrate IF-mediated AMPK signaling counteracts two key molecular drivers of RV failure. Thus, IF may be a novel treatment approach for RV dysfunction, a currently untreatable and lethal consequence of PAH.

**Highlights:**

- Intermittent fasting activates AMPK to restructure right ventricular mitochondrial and peroxisomal fatty acid fatty acid metabolism in two rodent models of PAH.
- Intermittent fasting prevents downregulation of multiple electron transport chain proteins in both monocrotaline and Sugen-hypoxia RVs.
- Pathological microtubule-mediated junctophilin-2 dysregulation and subsequent t-tubule remodeling is mitigated by intermittent fasting.
- Intermittent fasting suppresses the induction of both the canonical and peroxisomal ferroptosis pathways in RV failure.

## Introduction

IF is a powerful life-prolonging intervention in nondiseased states as numerous studies demonstrate IF confers survival benefits in multiple species and in several disease models(1,2). One of the numerous biological benefits of IF is advantageous metabolic reprogramming(1). *C. elegans* studies show AMPK signaling specifically underlies the lifespan-extending properties of IF via augmentation of fatty acid oxidation(3). In particular, IF prevents the age-related decline in mitochondria and peroxisomes, and thereby enhances lipid metabolism(3).

RV failure is the leading cause of death in PAH, and unfortunately there are no effective therapies that directly enhance RV function(4). Multiple laboratories have linked impaired fatty acid oxidation and altered lipid metabolism to RV dysfunction in PAH(5). Both preclinical models(6,7) and human data(8) demonstrate increased levels of toxic ceramides in the RV, which can be a consequence of disrupted mitochondrial fatty acid degradation(9). We recently showed IF combats lipotoxicity as there is an attenuation of ceramide accumulation in the RV of the monocrotaline (MCT) rat model of PAH(10). Thus, IF may enhance or even restructure RV lipid metabolism, but an in-depth molecular characterization of the effects of IF on RV mitochondrial and peroxisomal lipid handling and metabolizing enzymes is lacking.

In addition to AMPK’s well documented metabolic effects, AMPK also regulates microtubule dynamics via phosphorylation of the microtubule-associated proteins, microtubule associated protein-4 (MAP4)(11) and cytoplasmic linker protein-170 (CLIP170)(12). Phosphorylation of MAP4 and CLIP170 reduces their microtubule binding affinity, which ultimately suppresses their microtubule stabilizing properties. There is strong evidence that increased microtubule density/stability compromises cardiac function(13,14). We and others show microtubule remodeling results in disrupted transverse-tubule (t-tubule) structure and function via dysregulation of junctophilin-2(15,16). However, the impact of IF on MAP4 and CLIP170 phosphorylation states and the subsequent effects on microtubule regulation and t-tubule architecture have not been explored.

## Methods

An expansive methods section is available in the Supplemental Appendix. Male Sprague Dawley rats were randomized into control, MCT-induced PAH, and MCT with every other day fasting as we described(10). In addition, we randomized male Sprague Dawley rats to control, Sugen-hypoxia (SuHx) *ad lib* feeding, and Su-Hx every other day fasting. To be more translational, IF in SuHx rats was started after the return to normoxia, a timepoint when PH is established(17). We chose male rats due to their more severe RV phenotype as compared to females(18). Immunoblots analyzed AMPK activation and protein abundance. Electron microscopy evaluated RV mitochondrial morphology and cristae organization. Confocal microscopy determined peroxisomal density, microtubule remodeling, and t-tubule structure in RV cardiomyocytes. RV mitochondrial/peroxisomal enrichments were subjected to quantitative mass spectrometry and then the normalized values of protein abundance were correlated with tricuspid annular plane systolic excursion and Ees/Ea as determined by echocardiography and closed-chest pressure-volume loop analysis as we have previously described(7,10,19). ETC activity from frozen RV specimens was determined using Agilent XFe96 Seahorse apparatus.

## Results

### IF Activated AMPK, Modulated Mitochondrial and Peroxisomal Morphology, and Restructured Fatty Acid Metabolizing/Handling Enzymes Regulation in MCT and Su-Hx Rats

We first evaluated how IF modulated AMPK activation in the RV in the MCT model of RV failure. Immunoblots revealed MCT rats allowed to eat *ad libitum* (MCT-*ad lib*) had blunted AMPK signaling with lower levels of phosphorylated AMPK. However, IF activated AMPK as demonstrated by a significant increase in the levels of phosphorylated AMPK (pAMPK) and the ratio of pAMK to total AMPK as compared to MCT-*ad lib* animals (**Figure 1A and B**). Because AMPK regulates mitochondria and peroxisomal form and function(3), we used electron microscopy to evaluate mitochondrial morphology and cristae structure. There were no significant differences in mitochondrial length to width ratios when comparing the three groups, but IF partially restored mitochondrial cristae morphology (**Figure 1C**). Confocal microscopy analysis was used to determine how IF impacted peroxisomal density. When evaluating RV cardiomyocyte peroxisomes, we observed a significant increase in peroxisomal density in MCT-*ad lib* animals, but IF blunted this abnormal response (**Figure 1D**). To understand how these morphological changes were related to fatty acid metabolic protein regulation, we performed quantitative proteomics profiling of RV mitochondrial and peroxisomal enrichments. Hierarchical cluster analysis demonstrated MCT rats had a distinct mitochondrial fatty acid proteomic signature as compared to controls. There were higher levels of the Acyl-CoA Synthetase Long Chain (ACSL) family of proteins in MCT-*ad lib* RVs, but lower levels of nearly all proteins responsible for fatty acid degradation including both units of the trifunctional enzyme (HADHA and HADHB) and multiple acyl-CoA dehydrogenase (ACAD) family members (**Figure 1E**). However, IF prevented the upregulation of ACSL proteins and downregulation of HADHA/HADHB and multiple ACAD proteins (**Figure 1E**). When peroxisomal α- and β-oxidation proteins were profiled, we observed differences between controls and MCT-*ad lib*, which was again abrogated by IF (**Figure 1F**). In MCT-*ad lib* RVs, multiples enzymes responsible for the import, export, and degradation of lipid species were upregulated. Interestingly, not all peroxisomal fatty acid metabolizing enzymes were increased in MCT-*ad lib* RVs as phytanoyl-CoA 2-hydroxylase (PHYH), L-bifunctional enzyme (EHHADH), sterol carrier protein 2 (SCP2), and peroxisome 3-ketoacyl-CoA thiolase (ACAA1) had decreased abundance in MCT-*ad lib* animals (**Figure 1F**). These results suggested both mitochondrial and peroxisomal fatty acid metabolic pathways were reprogramed in RV dysfunction, and that IF partially negated these responses.

**Figure 1:**
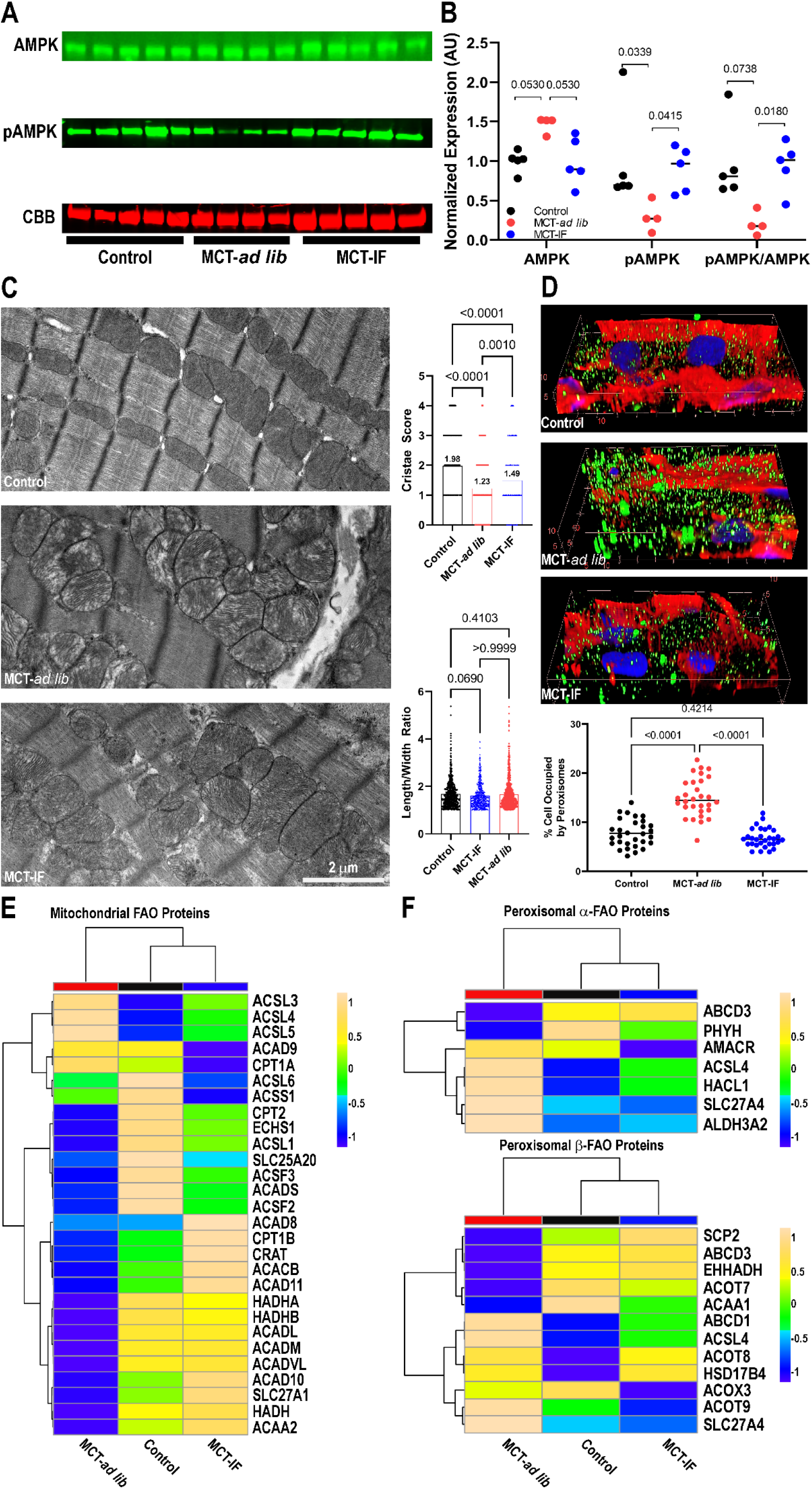
Intermittent fasting activated AMPK signaling and restructured mitochondrial morphology and peroxisomal density and lipid metabolism in MCT rats. (A) Representative Western blots of total AMPK and phosphorylated AMPK from RV extracts. CBB is Coomassie brilliant blue stained gel in the area of the myosin heavy chain. (B) Quantification of relative protein abundance. IF significantly increased pAMPK and pAMK/AMPK, consistent with an AMPK activating effect. *p*-values as determined by Kruskal-Wallis test with Dunn’s multiple comparisons test. (C) Representative electron micrographs of RV specimens depicting mitochondrial size and cristae morphology. Quantification of mitochondrial cristae morphology and length:width ratios. *p*-values as determined by Kruskal-Wallis with Dunn’s multiple comparisons test. (D) Representative three-dimensional reconstructions of confocal micrographs of RV cardiomyocytes stained with catalase antibody (green), wheat germ agglutinin (WGA) (red), and DAPI (blue) showing cardiomyocyte peroxisomes. IF prevented the increase in peroxisomal density. *p*-values as determined by one-way ANOVA with Tukey’s multiple comparisons test. (E) Hierarchical cluster analysis of mitochondrial fatty acid oxidation proteins. MCT-*ad lib* animals had reduced levels of multiple fatty acid oxidation proteins, which were mitigated by IF. (F) Hierarchical cluster analysis of peroxisomal α- and β-fatty acid oxidation proteins. IF peroxisomal fatty acid signature was comparable to controls and distinct from MCT-*ad lib*.

Next, we evaluated the effects of IF in SuHx rats with established PH. Consistent with our MCT results, IF activated RV AMPK and improved mitochondrial cristae structure and mitochondrial length to width ratio (**Figure 2 A-C**). Likewise, IF partially suppressed the increase in RV peroxisomal density (**Figure 2D**). Proteomics analysis revealed a very similar pattern of dysregulation of both mitochondrial and peroxisomal fatty acid metabolic/handling proteins in SuHx-*ad lib* RVs, which again was partially mitigated by IF (**Figure 2 E and F**).

**Figure 2:**
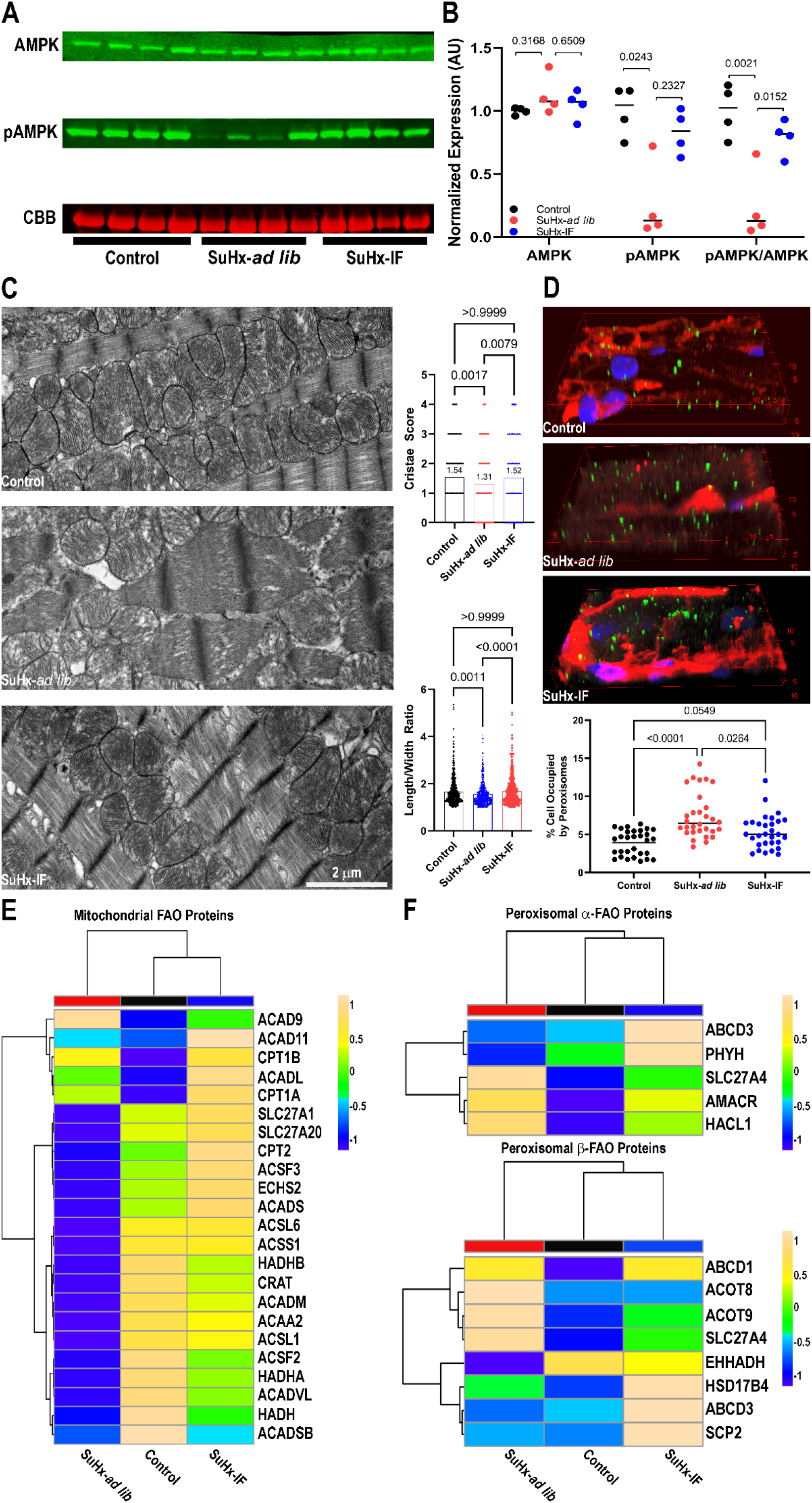
Intermittent fasting restored levels of phosphorylated AMPK and corrected mitochondrial architecture, peroxisomal density, and lipid metabolic proteins in Sugen-hypoxia rats. (A) Representative Western blots of total AMPK and phosphorylated AMPK from RV extracts. CBB is Coomassie brilliant blue stained gel in the area of the myosin heavy chain. (B) Quantification of relative protein abundance. IF significantly increased pAMPK and pAMK/AMPK, consistent with an AMPK activating effect. *p*-values as determined by Kruskal-Wallis test with Dunn’s multiple comparisons test. (C) Representative electron micrographs of RV specimens depicting mitochondrial size and cristae morphology. Quantification of mitochondrial cristae morphology and length:width ratio. *p*-values as determined by Kruskal-Wallis with Dunn’s multiple comparisons test. (D) Representative three-dimensional reconstructions of confocal micrographs of RV cardiomyocytes stained with catalase antibody (green), WGA (red), and DAPI (blue) showing cardiomyocyte peroxisomes. IF prevented the increase in peroxisomal density. *p*-values as determined by Kruskal-Wallis with Dunn’s multiple comparisons test. (E) Hierarchical cluster analysis of mitochondrial fatty acid oxidation proteins. SuHx-*ad lib* animals had reduced levels of multiple fatty acid oxidation proteins, which was mitigated by IF. (F) Hierarchical cluster analysis of peroxisomal α- and β-fatty acid oxidation proteins. IF peroxisomal fatty acid signature was comparable to controls and distinct from SuHx-*ad lib*.

### IF Combatted Dysregulation of Electron Transport Chain Proteins in Both Rodent Models

In order to harness the energetic potential of fatty acid oxidation, the electron transport chain needs to convert NADH and FADH_2_ into ATP(20). Therefore, we examined the effects of IF on electron transport chain protein abundance and function. MCT-*ad lib* RVs had a distinct proteomic signature as compared to controls when each electron transport chain complex subunit was profiled. Hierarchical cluster analyses of all five electron transport chain complexes showed IF animals clustered with controls as IF prevented the downregulation of multiple electron transport chain subunit proteins that was observed in MCT-*ad lib* rats (**Figure 3A-E**). Next, we quantified electron transport chain activity using a recently developed Seahorse-based assay(21). Complex I and Complex II oxygen consumption rates were reduced in MCT-*ad lib* RVs, but IF normalized the oxygen consumption rates of both complexes (**Figure 3F and G**). Complex IV activity was not significantly different between the three groups, however the sum of all three complexes was depressed in MCT-*ad lib* but restored by IF (**Figure 3H and I**). Thus, these data demonstrated IF corrected RV electron transport chain protein regulation and activity.

**Figure 3:**
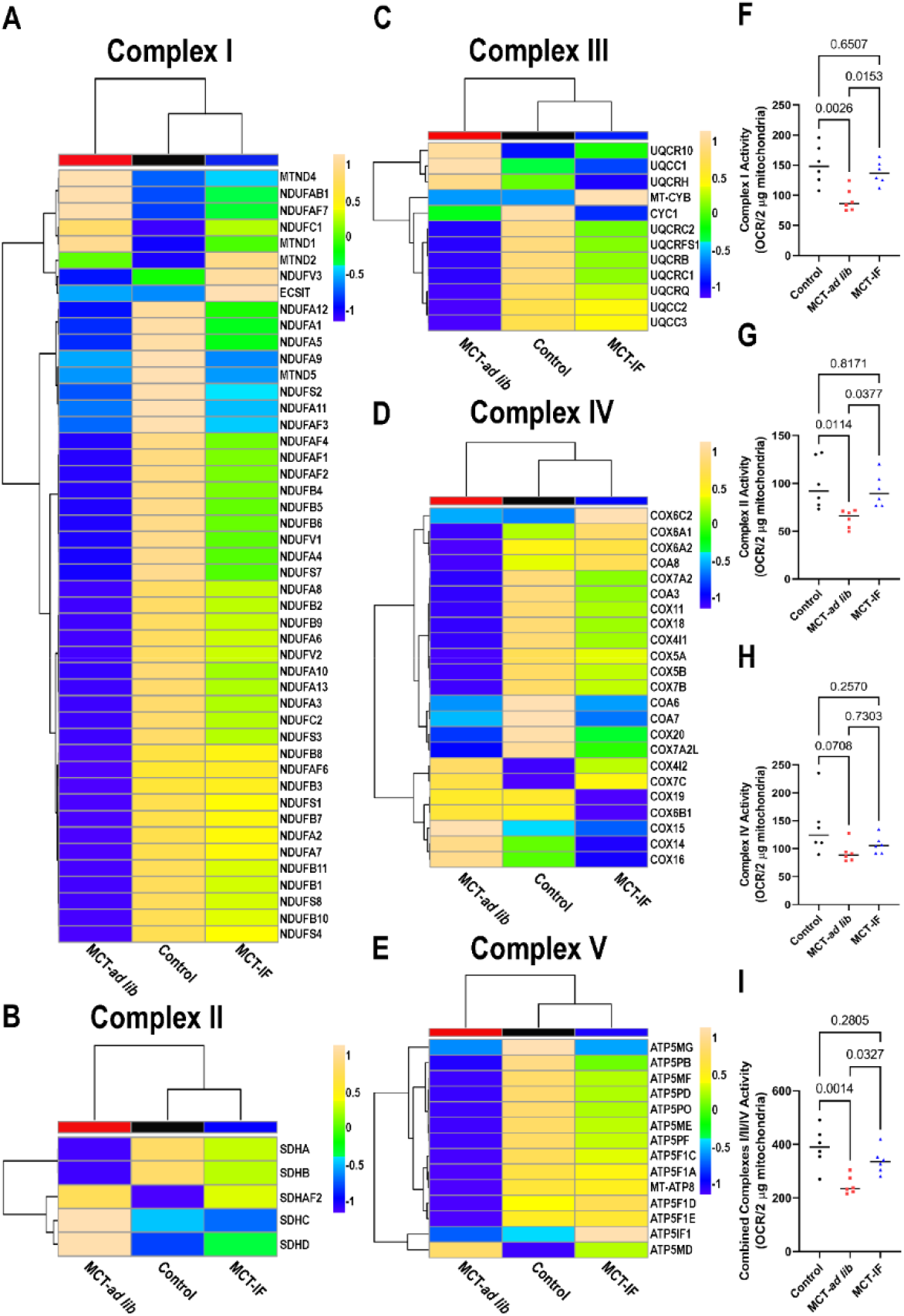
Intermittent fasting prevented dysregulation of electron transport chain protein subunits and enhanced electron transport chain function in MCT rats. Hierarchical cluster analyses of proteomic-derived abundance of proteins in Complex I (A), Complex II (B), Complex III (C), Complex IV (D), and Complex V (E). Intermittent fasting prevents dysregulation of all ETC complexes. Quantification of Complex I (F), Complex II (G), Complex IV (H), and combined Complex I, II, and IV (I) activity of frozen RV extracts. Intermittent fasting augmented electron transport chain function. *p*-values as determined by one-way ANOVA with Tukey’s multiple comparisons test.

When we evaluated the effects of IF on electron transport chain protein regulation in SuHx rats, we again observed very similar results as those obtained in the MCT rats. Numerous subunits in Complex I (**Figure 4 A**), Complex II (**Figure 4B**), Complex III (**Figure 4C**), Complex IV (**Figure 4D**), and Complex V (**Figure 4E**) were downregulated in SuHx-*ad lib* RVs, but IF mitigated these changes. In fact, hierarchical cluster analysis demonstrated SuHx-IF animals clustered with controls in all five complexes.

**Figure 4:**
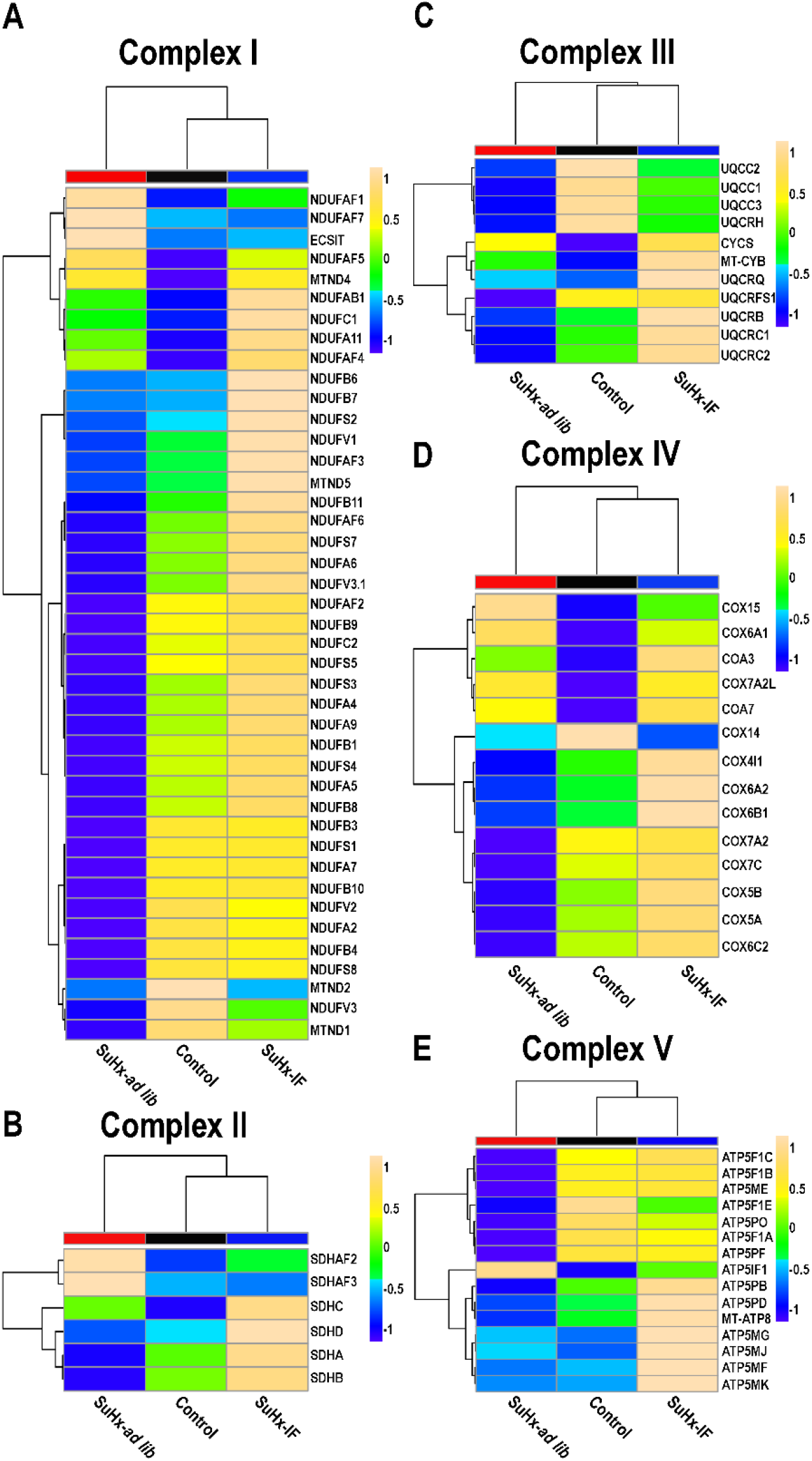
Intermittent fasting combatted downregulation of electron transport chain protein subunits in Sugen-hypoxia rats. Hierarchical cluster analyses of proteomic-derived abundance of proteins in Complex I (A), Complex II (B), Complex III (C), Complex IV (D), and Complex V (E).

### Correlational Heatmapping Identified Associations Between RV Function and Fatty Acid Metabolism and Electron Transport Chain Proteins

Next, we used correlational heatmapping to interrogate the relationships between tricuspid plane annular systolic excursion (TAPSE), pressure-volume loop derived right ventricular-pulmonary artery coupling (Ees/Ea), and fatty acid oxidation and electron transport chain protein abundance. We observed significant and positive correlations between multiple mitochondrial fatty acid oxidation proteins with TAPSE and Ees/Ea (**Figure 5A-D**). However, ACSL4/5 protein abundance was inversely associated with RV function (**Figure 5A-D**). Peroxisomal fatty acid proteins also significantly correlated with RV function. Interestingly, enzymes responsible for fatty acid degradation were associated with better RV function, but proteins that modulate lipid uptake were negatively associated with RV function (**Figure 5E-H**). Finally, we showed positive correlations between multiple electron transport chain proteins in all five complexes and RV function (**Figure 5I-L**).

**Figure 5:**
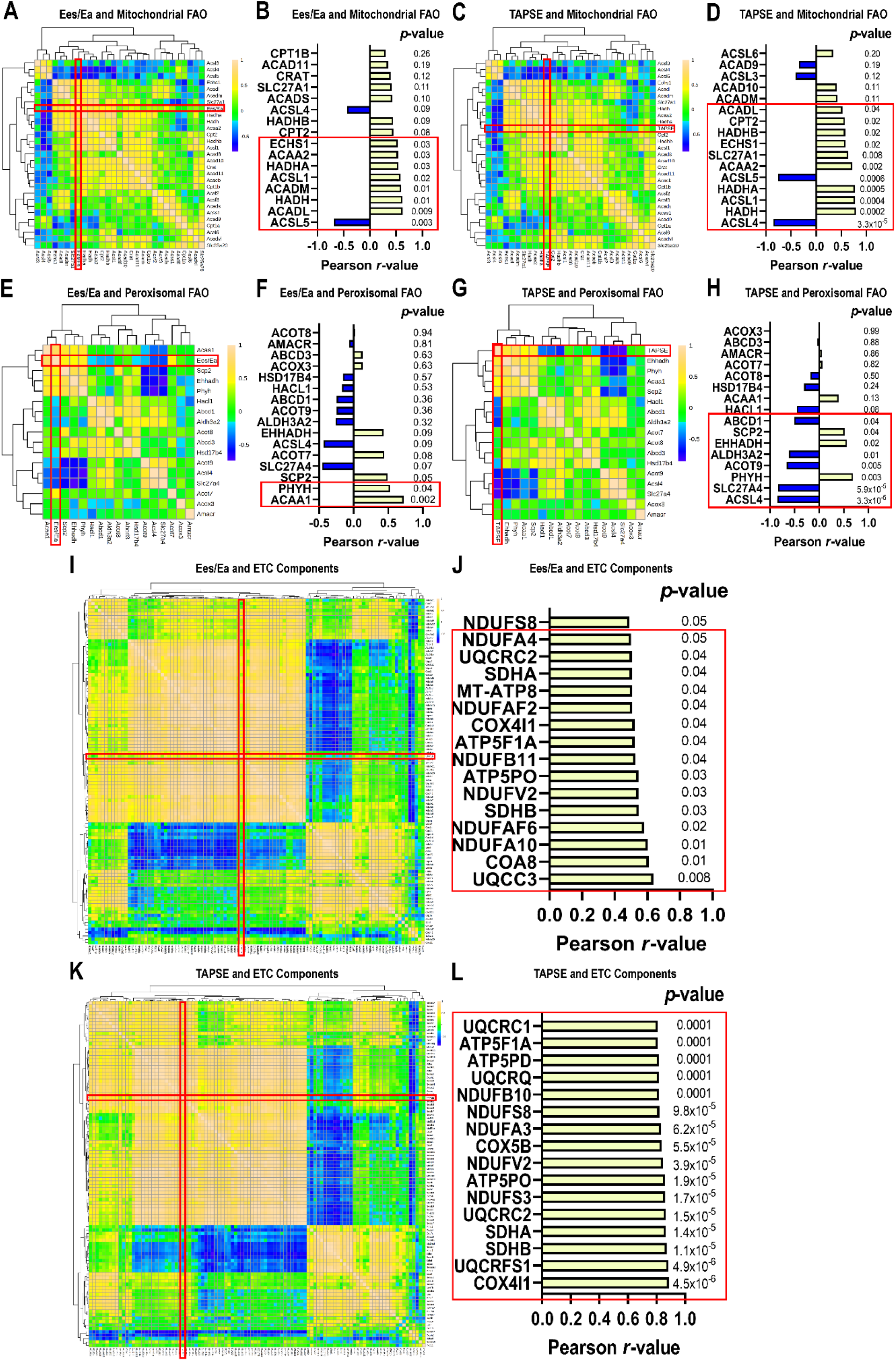
Correlational heatmapping identified relationships between fatty acid oxidation, electron transport chain proteins, and RV function in MCT rats. (A) Correlational heat map of mitochondrial fatty acid oxidation proteins and Ees/Ea. (B) *r*-value and *p*-value for each protein’s relationship with Ees/Ea.. (C) Correlational heat map of mitochondrial fatty acid oxidation proteins and TAPSE. (D) *r*-value and *p*-value for each protein’s relationship with TAPSE. Red box highlights significant relationships. (E) Correlational heat map of peroxisomal fatty acid oxidation proteins and Ees/Ea. (F) *r*-value and *p*-value for each protein’s relationship with Ees/Ea. (G) Correlational heat map of peroxisomal fatty acid oxidation proteins and TAPSE. (H) *r*-value and *p*-value for each protein’s relationship with TAPSE. (I) Correlational heat map of electron transport chain proteins and Ees/Ea. (J) *r*-value and *p*-value for each protein’s relationship with Ees/Ea. (K) Correlational heat map of electron transport chain proteins and TAPSE. (L) *r*-value and *p*-value for each protein’s relationship with TAPSE. In all figures, red boxes highlight significant relationships.

When we performed correlational heatmapping analysis in the SuHx rodents, our results slightly diverged from the MCT results. Again, many mitochondrial fatty acid metabolizing proteins were positively associated with RV function (**Figure 6A-D**), but many ACSL proteins were not detected in SuHx proteomics experiments so that negative associated was absent. When peroxisomal fatty acid proteins were probed, we found a negative association between fatty acid import proteins and RV function and a nonsignificant but positive relationship between EHHADH and RV function (**Figure 6E-H**). The most divergent observations were found when we correlated ETC protein abundance and RV function. In SuHx experiments, again there were multiple proteins positively associated with RV function, but many were negatively associated with RV function (**Figure 6I-L**).

**Figure 6:**
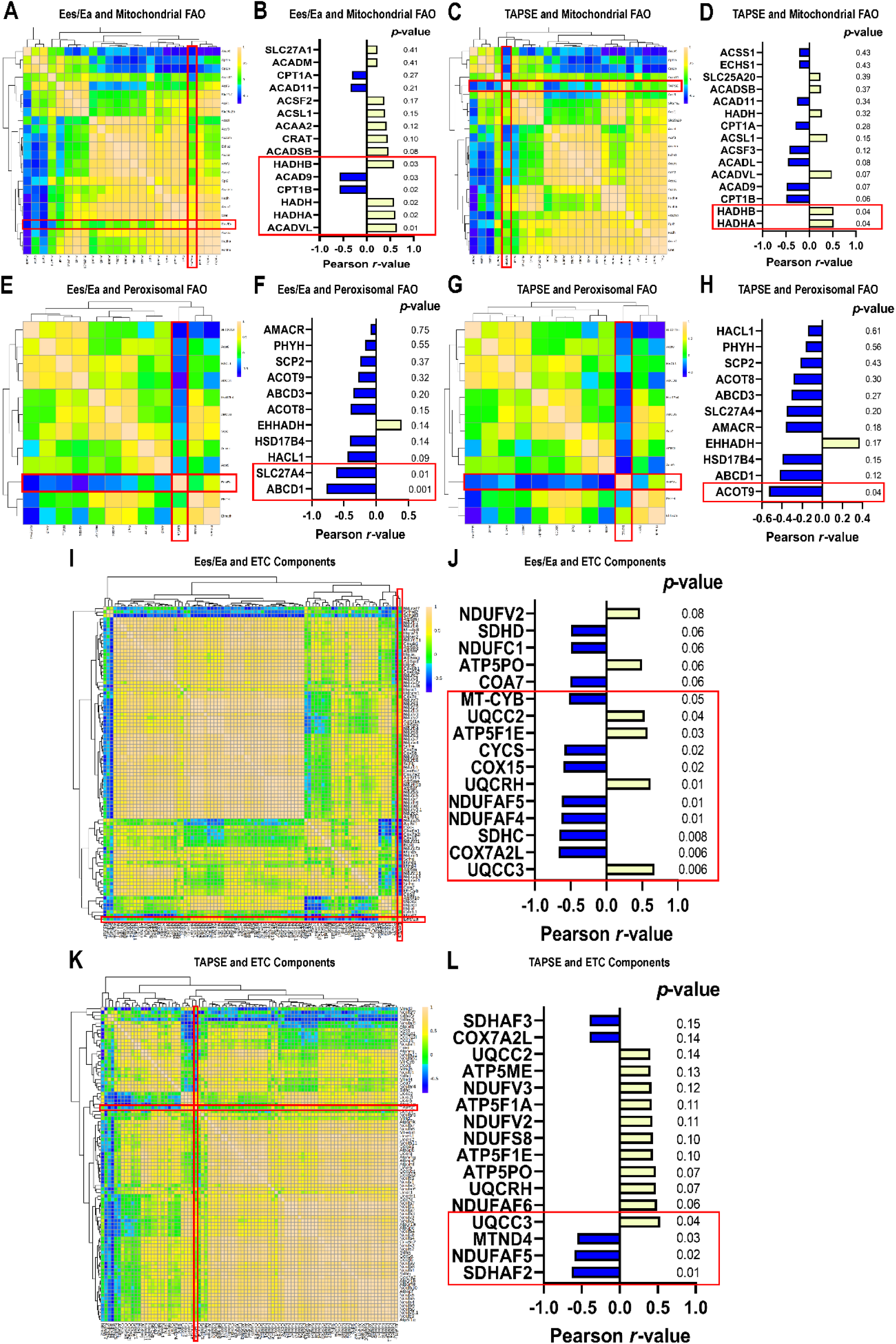
Correlational heatmapping identified comparable relationships between fatty acid oxidation, electron transport chain proteins, and RV function in Sugen-hypoxia rats. (A) Correlational heat map of mitochondrial fatty acid oxidation proteins and Ees/Ea. (B) *r*-value and *p*-value for each protein’s relationship with Ees/Ea. (C) Correlational heat map of mitochondrial fatty acid oxidation proteins and TAPSE. (D) *r*-value and *p*-value for each protein’s relationship with TAPSE. (E) Correlational heat map of peroxisomal fatty acid oxidation proteins and Ees/Ea. (F) *r*-value and *p*-value for each protein’s relationship with Ees/Ea. (G) Correlational heat map of peroxisomal fatty acid oxidation proteins and TAPSE. (H) *r*-value and *p*-value for each protein’s relationship with TAPSE. (I) Correlational heat map of electron transport chain proteins and Ees/Ea. (J) *r*-value and *p*-value for each protein’s relationship with Ees/Ea. (K) Correlational heat map of electron transport chain proteins and TAPSE. (L) *r*-value and *p*-value for each protein’s relationship with TAPSE. In all figures, red boxes highlight significant relationships.

### Intermittent Fasting Suppressed Pathological Microtubule Remodeling and Corrected T-tubule Architecture

Then, we evaluated how AMPK activation modulated the microtubule cytoskeleton in RV failure. Western blot analysis revealed higher levels of the microtubule subunit proteins, α- and β-tubulin, in MCT-*ad lib* RVs, which was prevented by IF (**Figure 7A and B**). To determine if altered phosphorylation patterns of MAP4 and CLIP170 contributed to normalization of tubulin regulation, we probed total and phosphorylated levels of each protein in RV extracts. Both total CLIP170 and MAP4 were elevated in MCT-*ad lib* RVs, which IF combatted (**Figure 7A and B**). The three groups showed no differences in the ratio of phosphorylated CLIP170/total CLIP170, but the pMAP4/MAP4 ratio in the MCT-*ad lib* RVs was significantly depressed, which IF partially, but nonsignificantly corrected (**Figure 7A and B**). Then, we used confocal microscopy to quantify microtubule structure in RV cardiomyocytes. MCT-*ad lib* microtubule lattice density was significantly increased as compared to controls, but IF blunted microtubule proliferation (**Figure 7C and D**). Finally, we evaluated the downstream effects of microtubule dysregulation by probing junctophilin-2 protein abundance and t-tubule structure. Consistent with our previous work(15,19), junctophilin-2 levels were lower (**Figure 7A and B**) and t-tubule morphology was deranged in MCT-*ad lib* RV cardiomyocytes (**Figure 7 E and F**). However, in MCT-IF rats, junctophilin-2 levels were normalized (**Figure 7 A and B**) and t-tubule architecture was restored (**Figure 7E and F**).

**Figure 7:**
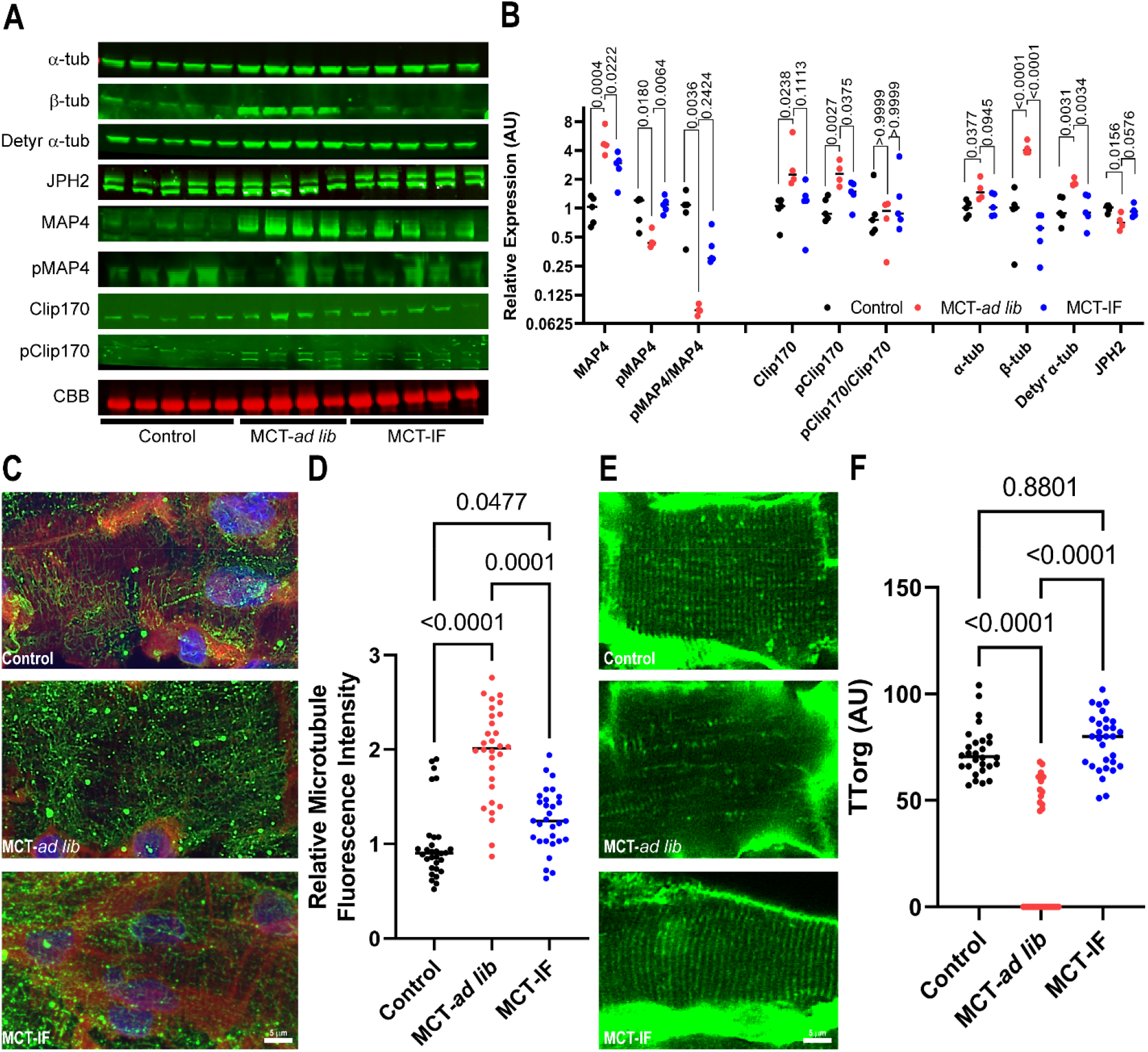
Intermittent fasting mitigated pathological microtubule remodeling, which restored junctophilin-2 expression, and rescued t-tubule structure in RV cardiomyocytes of MCT rats. (A) Western blots and subsequent quantification (B) of tubulins and microtubule regulating proteins. (C) Representative confocal micrographs of microtubules in RV cardiomyocytes. (D) Quantification of relative microtubule density. IF mitigates increased microtubule density. *p*-values as determined by Kruskal-Wallis with Dunn’s multiple comparisons test. (E) Representative images of RV cardiomyocyte t-tubules. (F) Quantification of t-tubule organization. *p*-values as determined by Kruskal-Wallis with Dunn’s multiple comparisons test.

Finally, we examined the effects of IF on microtubule and t-tubule regulation in SuHx rats. Although not as robust as in MCT animals, we found IF prevented upregulation of multiple tubulin subunits, elevated junctophilin-2 levels, and increased the ratio of pMAP4/MAP4 (**Figure 8 A and B**). Furthermore, IF combated t-tubule structural abnormalities (**Figure 8 C and D**).

**Figure 8:**
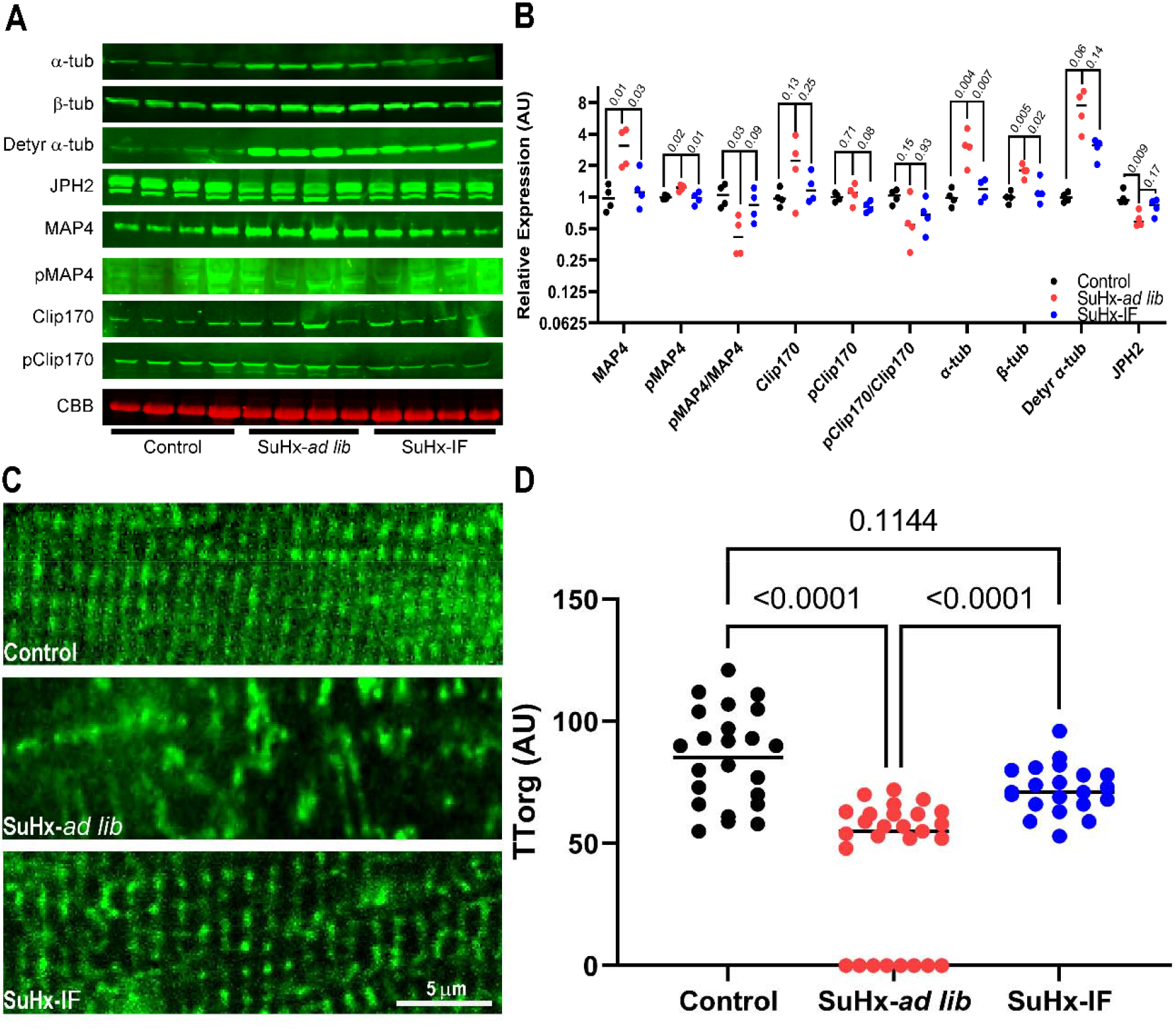
Intermittent fasting partially prevented microtubule dysregulation and improved t-tubule morphology in the RV of Sugen-hypoxia rats. (A) Western blots and subsequent quantification (B) of tubulins, microtubule regulating proteins, and junctophilin-2. (C) Representative images of RV cardiomyocyte t-tubules. (D) Quantification of t-tubule organization. *p*-values as determined by Kruskal-Wallis with Dunn’s multiple comparisons test.

## Discussion

In summary, we show IF prevents dysregulation of both mitochondrial and peroxisomal fatty acid metabolism, restores mitochondrial cristae structure, suppresses peroxisomal proliferation, counteracts dysregulation of multiple ETC subunits, and combats microtubule-mediated t-tubule derangements in two rodent models of PAH-mediated RV failure. In general, the effects of IF were slightly less robust in the SuHx rats as what was observed in MCT rats, and this may be due to differences in the two models and the timing of the IF intervention. However, the consistent observation that IF activates AMPK which restructures fatty acid metabolism, microtubules, and ETC regulation suggests it may have promise as a nonpharmacological intervention in patients with RV failure.

In our correlational analyses in the MCT experiments, we observed an inverse relationship between ACSL4/5 and RV function, which requires further discussion and exploration. ACSL4/5 regulate ferroptosis(22,23), a nonapoptotic form of cell death linked to alterations in lipid metabolism(24). To understand how ferroptosis may modulate RV function in PAH, we probed ferroptotic protein regulation and the relationships between ferroptosis proteins and TAPSE and Eea/Ea. Hierarchical cluster analysis showed MCT-*ad lib* had higher levels of pro-ferroptotic proteins than control animals, and the overall pattern of ferroptotic-regulating proteins was distinct (**Supplemental Figure 1A**). However, IF animals had a ferroptosis molecular signature that was similar to control with less upregulation of pro-ferroptotic enzymes (**Supplemental Figure 1A**). When performing correlational heatmapping, we found pro-ferroptotic proteins were negatively associated with RV function with one exception: Raf kinase inhibitor protein (RKIP/PEBP1) (**Supplemental Figure 1**).

Consistent with our findings, PEBP1 directly augments cardiac contractility via modulation of beta-adrenergic signaling(25), and perhaps the positive inotropic effect of PEBP1 is more important than its ferroptosis function in diseased states. We also observed a pro-ferroptotic signature in the RV of SuHx rats, and there was a trend for a reduction in pro-ferroptotic proteins in the SuHx RV specimens (**Supplemental Figure 2**). However, we did not see as a profound effect with IF in SuHx rats as hierarchical cluster analysis revealed SuHx-IF had a ferroptotic signature that was more similar to SuHx-*ad lib* than control (**Supplemental Figure 2**). The less potent effect may be due to initiation of IF at a later timepoint in the disease process and differences in protein detection as the SuHx proteomics analysis did not detect as many ferroptosis proteins. Nonetheless, these results suggest another beneficial effect of IF is antagonism of ferroptosis. It is very possible the anti-ferroptotic effect of IF is also mediated by AMPK because AMPK suppresses ferroptosis in cancer cells and in renal ischemia-reperfusion by reprogramming lipid metabolism(26).

To the best of our knowledge, this study is the first to interrogate the effects of IF on ferroptosis protein regulation and cardiac function. However, our finding that a pro-ferroptosis state is associated with depressed cardiac contractility is congruent with results by other groups. First, small molecule inhibition of ferroptosis combats both doxorubicin and ischemia-reperfusion mediated left ventricular dysfunction(27). Interestingly, the dampening of ferroptosis restores Complex I and II activity(27), which is consistent with our observation that conditions of heightened ferroptosis result in impaired Complex I and II function. Second, suppression of ferroptosis via glutathione peroxidase 4, a protein that reverses lipid peroxidation and blunts ferroptosis induction, overexpression mitigates doxorubicin cardiomyopathy(28). Furthermore, mice with reduced glutathione peroxidase 4 (GPx4) expression have worse cardiac function after doxorubin exposure(28). The combination of these publications and our data show excess ferroptosis promotes cardiac dysfunction. Certainly, future studies examining the effects of ferroptosis inhibition in RV failure due to PAH would be helpful to determine if this pathway may be a novel therapeutic target.

Our current findings of peroxisomal remodeling in the setting of RV dysfunction agree with our previous work(7), but the consequences of these observations remain unknown. However, the association between increased peroxisomal abundance and RV dysfunction along with the fact that many peroxisomal proteins are inversely correlated with RV function implies peroxisomal reprogramming may be maladaptive. A potential mechanism for this hypothesis could also be ferroptosis induction, as a large CRISPR-based screen identified multiple peroxisomal proteins have the capacity to modulate ferroptosis sensitivity(29). A series of well executed experiments revealed the synthesis of polyunsaturated ether phospholipids through the peroxisomal enzymes fatty acyl-CoA reductase 1 (FAR1), glyceronephosphate *O*-acyltransferase (GNPAT), and alkylglycerone phosphate synthase (AGPS) and the endoplasmic reticulum enzyme 1-acylglycerol-3-phosphate *O*-acyltransferase 3 (AGPAT3) drives the peroxisomal-ferroptotic pathway(29). When we profiled these particular proteins in our proteomics experiments, we found increased abundance of all enzymes in MCT-*ad lib* RV (**Supplemental Figure 3**). IF partially suppressed the upregulation of all of these proteins, and hierarchical cluster analysis demonstrated MCT-IF rats had a similar peroxisomal-ferroptotic signature to controls animals (**Supplemental Figure 3**). Interestingly, all peroxisomal-ferroptotic pathway enzymes were inversely associated with RV function, and in particular, AGPS had the strongest association with RV dysfunction (**Supplemental Figure 3**). Again, we observed very consistent findings in our SuHx experiments as some peroxisomal ferroptotic proteins were upregulated in the SuHx RV and the changes were blunted by IF (**Supplemental Figure 4**), which further supports the hypothesis that peroxisomal ferroptosis could negatively impact cardiac function. In conclusion, our study raises the possibility that increased peroxisomal density and the associated changes in enzymatic regulation may have deleterious effects on RV function via a newly described peroxisomal-ferroptotic pathway.

When evaluating the effects of IF, it is important to acknowledge that caloric restriction could underlie the benefits observed in this study. However, there are multiple lines of evidence that suggest IF is more effective than caloric restriction at modulating metabolism and extending lifespan. First, the survival benefit conferred by IF in MCT rats cannot be explained by caloric restriction alone because MCT-IF rats ate as much food as the MCT-*ad lib* rats in our previous study(10). However, in the Sugen-hypoxia rat experiments, SuHx-IF rats ate less total food than the SuHx-*ad lib* rodents, although not statistically significant, (**Supplemental Figure 5**) so caloric restriction may have played a role. In addition to our findings, Pak *et al* specifically showed fasting, and not caloric restriction, underlies the heightened insulin sensitivity and greater utilization of fatty acids as fuel in aged mice(30). Moreover, Acosta-Rodriguez *et al* demonstrated daily fasting properly aligned with the circadian rhythm extends lifespan beyond that of caloric restriction (35% versus 10% increase)(31). When beginning to consider a clinical trial of IF in PAH patients, integrating the circadian timing of fasting may be key for successful translation. Supporting this hypothesis, a recent clinical trial demonstrated early time restricted eating produces greater weight loss and drops diastolic blood pressure more than traditional caloric restriction(32). Clearly, there is a complex interplay between caloric restriction, fasting, and circadian rhythm, and it is likely that all three of these key variables will need to be accounted for when attempting to harness the beneficial effects of IF for RV failure.

Our manuscript has important limitations that we must acknowledge. First, IF has pleiotropic effects including restoring circadian rhythm, restructuring the gut microbiome, and systemic anti-inflammatory effects(1,2,33). Therefore, it is very unlikely that the modulation of RV lipid metabolism and microtubule density are the only molecular mechanisms underlying improved RV function. Second, our proteomics experiments were conducted using whole RV free wall specimens and thus nonmuscle cells may contribute to some of the findings. However, we did not detect significant differences in tubulin regulation or AMPK activation in RV fibroblasts with IF (**Supplemental Figure 6**). We were unable to calculate Ees/Ea in all animals during catheterization (*n*=1 MCT-*ad lib, n*=1 MCT-IF, *n*=1 SuHx-IF, and *n*=1 SuHx-*ad lib*), which may have explained where some of the correlational relationships were less significant than those observed with TAPSE. Finally, in the SuHx proteomics experiments, two of the SuHx-IF samples were inadvertently mixed together during processing for proteomics analysis. The two samples were run in duplicate but only was sample was included in the hierarchical cluster analysis. Neither samples were used for correlational heatmapping analysis, which reduced our sample size and may have contributed to some of the differences between MCT and SuHx animals in our correlational heat mapping analysis.

## Acknowledgements

The authors thank Drs. LeeAnn Higgins and Todd Markowski of the University of Minnesota Center for Mass Spectrometry and Proteomics for assistance with our proteomics experiments, Dr. Keita Uchida for sharing his antigen retrieval protocol for cardiomyocyte microtubule imaging, Mr. Trace Christensen and the Mayo Microscopy and Cell Analysis Core for experimental and technical support, Ms. Bridget Nieto and Dr. Michael Cypress for experimental and technical support, and Dr. Yasunori Shintani for generously sharing the phospho-CLIP170 antibody.

## Perspectives

Competency in Medical Knowledge: Although RV failure is the leading cause of death in PAH, we still do not have therapies that directly improve RV dysfunction. Here, we show impaired fatty acid oxidation and microtubule proliferation are associated with RV dysfunction in two models of rodent PAH. Counteracting these two biological processes via intermittent fasting could improve RV function and hopefully extend lifespan in PAH.

Translational Outlook: Intermittent fasting may be a nonpharmacological means to improve RV function in PAH, and potentially synergize with current therapies. Further human studies are needed to understand if PAH patients receive any benefits from intermittent fasting.

## Online Methods

### Animal Studies

Male Sprague Dawley rats, purchased from Charles River Laboratories, were randomized into three experimental groups: phosphate buffered saline (PBS) injected (control), monocrotaline-*ad libitum*, and monocrotaline-IF. All monocrotaline rats received a single subcutaneous injection of monocrotaline (60 mg/kg) (Sigma-Aldrich, St. Louis, MO). The monocrotaline-IF group began every other day fasting the day following monocrotaline injection. For Sugen-hypoxia experiments, male Sprague Dawley rats received a single subcutaneous injection of Sugen 5416 (20 mg/kg) (Cayman Chemical, Ann Arbor, MI) and were exposed to 10% oxygen for three weeks, returned to normoxia, and randomized into two experimental group: SuHx-*ad libitum*, and SuHx-IF. The SuHx-IF group began every other day fasting the day following SuHx treatment. Control group received a diluent injection and were maintained in normoxia until the end of study. All animal studies were approved by the University of Minnesota Institutional Animal Care and Use Committee.

### Western blot

Western blots were performed by probing a total of 25 μg of RV protein extract using the Odyssey Infrared Imaging system (Lincoln, NE) as previously described(15). Post transfer SDS-PAGE gels were stained with Coomassie brilliant blue (CBB) and imaged at the 700-nm wavelength on the Odyssey Imaging System with the band corresponding to the myosin heavy chain used as the loading control(15,34). For the fibroblast western blots, vimentin was used as the loading control instead of the CBB post transfer gel.

### Electron Microscopy

RV free wall tissue was placed into fixative (4% paraformaldehyde + 1% glutaraldehyde in 0.1M phosphate buffered, pH 7.2 (PB)). After fixation, tissue was washed with PB, stained with 1% osmium tetroxide, washed in H2O, stained in 2% uranyl acetate, washed in H_2_O, dehydrated through a graded series of ethanol and acetone and embedded in Embed 812 resin. Following a 24 hour polymerization at 60°C, 0.1 µM ultrathin sections were prepared and post-stained with lead citrate. Micrographs were acquired using a JEOL 1400 Plus transmission electron microscope (JEOL, Inc., Peabody, MA) at 80 kV equipped with a Gatan Orius camera (Gatan, Inc., Warrendale, PA).

### Confocal microscopy

RV cardiomyocyte bundles from formalin fixed RV free wall specimens were manually dissected using a dissecting microscope. Bundles were subjected to antigen retrieval using Reveal Decloaker solution (Biocare Medical) for 45 minutes at 95^0^C. Samples were allowed to cool to room temperature and then incubated in 1% Triton X-100 in PBS for 10 minutes. Samples were blocked in 5% goat serum in PBS for ten minutes. Primary antibodies diluted in 5% goat serum in PBS were incubated with bundles at 4^0^C for 48-72 hours. Bundles were washed/blocked in 5% goat serum and then incubated with secondary antibody and Wheat Germ Agglutinin Alexa Fluor-633 conjugate (ThermoFisher) at 37^0^C for 30 minutes. Samples were washed with PBS, treated with an autofluorescence quenching kit (Vector Laboratories), and then embedded in Antifade containing DAPI (Vector Laboratories). Z-stacks of cardiomyocyte bundles were obtained with an Olympus FV1000 BX2 upright confocal microscope (Tokyo, Japan) at the University of Minnesota Imaging Center.

### Quantification of mitochondrial structure

Mitochondrial gross morphology was quantified using a length to width ratio obtained from FIJI). Mitochondrial cristae structure was scored based on density percent of mitochondrial occupied by cristae and as previously described(35). All images were blindly analyzed by FK.

### Quantification of peroxisomal density

Cardiomyocyte bundles were stained for peroxisomes with a catalase antibody and density was determined using the threshold plug-in on FIJI (Bethseda, MD). Images were blindly analyzed by FK.

### Assessment of microtubule density

Cardiomyocyte bundles were stained with a β-tubulin antibody and then microtubule fluorescence intensity normalized to area was determined using FIJI (Bethseda, MD). Images were blindly analyzed by FK.

### T-tubule morphological analysis

WGA staining of cardiomyocyte bundles was used to delineate t-tubules. T-tubule regularity was quantified using the TT_Power_ plug-in on FIJI. FK blindly analyzed the images.

### Quantitative proteomics of mitochondrial/peroxisomal enrichments

RV mitocondrial/peroxisomal enrichment samples (*n*=5 control, *n*=5 MCT-*ad lib*, and *n*=6 MCT-IF) and (*n*=4 control, *n*=5 SuHx-*ad lib*, and *n*=6 SuHx-IF) were isolated with a mitochondrial isolation kit (Abcam). Then, the mitochondrial/peroxisome pellet fraction was resuspended in lysis buffer and supplemented with protease inhibitor (ThermoFisher). The samples were briefly sonicated with a Branson Digital Sonifier 250 (Branson Ultrasonics) and then underwent pressure cycling (alternating 35 KPSI for 20 seconds and 0 KPSI for 10 seconds for 60 cycles at 37°C). Protein concentration was determined by Bradford assay. A 25 µg aliquot of each sample was diluted in half with lysis buffer and then this sample was then diluted 5-fold with water. Trypsin (Promega) was added in a 1:40 ratio and samples were incubated overnight at 37°C and after incubation, frozen at -80°C and vacuum dried. Each sample was cleaned with a 1 cc Waters Oasis MCX cartridge (Waters Corporation), eluates were vacuum dried and resuspended in 100 µl of 0.1 M triethylammonium bicarbonate, pH 8.5 to a final concentration of 1 µg/µl. For each channel in the TMT16plex, a 20 µg aliquot of the sample was made and labeled with 0.2 mg TMT16plex. Isobaric Label Reagent (Thermo Scientific). After TMT labeling, all samples were multiplexed together.

We reconstituted the dried peptide fractions in 94.9:5:0.1, H2O:acetonitrile (ACN):formic acid (FA) (load solvent) and analyzed ∼500 nanograms of each fraction by capillary LC-MS with a Thermo Fisher Scientific, Inc Dionex UltiMate 3000 RSLCnano system on-line with an Orbitrap Eclipse mass spectrometer (Thermo Fisher Scientific) with FAIMS (high-field asymmetric waveform ion mobility) separation. We injected peptides directly in load solvent and separated peptides a flowrate of 315 nl/minute on a self-packed C18 column (Dr. Maisch GmbH ReproSil-PUR 1.9 um 120 Å C18aq, 100 um ID x 30 cm length) at 55 °C with the following gradient elution profile: from 0 – 2 minutes solvent B was held at 5%, from 2 – 2.5 minutes solvent B was ramped from 5 to 8%, from 2.5 – 135 minutes solvent B was ramped from 8 – 21%, from 135 – 180 minutes solvent B was ramped from 21 – 34%, from 180 – 182 minutes solvent B was ramped to 90% and held until 188 minutes, then at 190 minutes solvent B was changed to 5% B for for 7 minutes; solvent A was 0.1% formic acid in water and solvent B was 0.1% formic acid in ACN. The FAIMS total carrier gas was 4.6 and cooling gas setting was 5.0 L/min and the inner and outer electrodes were set to 100 °C. We scanned the CV (compensation voltage) at -45, 60 and 75 for 1 second each with a data dependent acquisition method. We employed the following MS parameters: ESI voltage 2.1 kV, ion transfer tube 275 °C; no internal calibration; Orbitrap MS1 scan 120k resolution in profile mode from 400 – 1400 *m/z* with 50 msec injection time; 100% (4E5) automatic gain control (AGC); HCD activation for precursors with 2 – 6 charges above 2.5E4 counts; MIPS (monoisotopic peak determination) set to Peptide mode; MS2 settings (all CV’s) were: 0.7 Da quadrupole isolation window, 38% collision energy, Orbitrap detection with 50K resolution at 200 *m/z*, first mass fixed at 110 *m/z*, 150 msec max injection time, 250% (1.25E5) AGC and 45 sec dynamic exclusion duration with +/- 10 ppm mass tolerance, dynamic exclusion list was not shared among CV’s and dependent scan on single charge state per precursor was set to False.

We processed peptide tandem MS using Sequest (Thermo Fisher Scientific, Inc) in Scaffold version 5 software. The rat Universal Proteome (UP000002494) protein sequence database was downloaded from UniProt (www.uniprot.org/) and merged with a common lab contaminant protein database (http://www.thegpm.org/cRAP/index.html). We applied the precursor mass recalibration node with the rat UP database, precursor mass tolerance 20 ppm, product ion tolerance 0.1 Da, dynamic mass TMTpro (304.2071 *m/z*) and fixed carbamidomethyl (CAM) modification of cysteine 57.0215 *m/z*. The Sequest (PMID: 24226387) database search parameters were: enzyme trypsin full specificity, 2 missed cleave sites; precursor tolerance 15 ppm, fragment ion tolerance was 0.08 Da. We set the dynamic modifications for TMTpro on lysine and peptide N-terminus, oxidation of methionine, pyroglutamic acid conversion of peptide N-terminal glutamine, deamidation of asparagine and glutamine. Dynamic protein N-terminal modifications were acetylation, M (methionine) loss (−131.040 Da) and M-loss + acetylation (−89.030 Da). We specified CAM cysteine as a fixed modification.

We used Scaffold version 5 for TMT-based protein quantification for the MCT experiments with the following parameters: unique and razor peptides were included, shared peptides were excluded, impurity corrections were applied, co-isolation threshold maximum was 50%, normalization was performed on the total peptide amount; protein ratio calculations were performed using pairwise ratio-based mode while applying a 1% protein and peptide False Discovery Rate (FDR). Hypothesis testing using the Benjamini-Hochberg false discovery rate procedure to control for errors associated with multiple hypothesis tests in Scaffold.

We used Proteome Discoverer version 3.0 for TMT-based protein quantification for the Su-Hx proteomics experiments with the same parameters as above.

### Assessment of electron transport chain activity

Mitochondrial respiration was measured using frozen tissues as described in Acin-Perez *et. al*(21). Briefly, snap-frozen RV tissue, stored at -80, was place in a 2 mL Kontes glass homogenizer with 200ul of MAS buffer then minced with a scissors. The tissue was homogenized with 20 strokes of pestle B (0.0005 inch clearance). Based on BCA assay (Thermo Scientific) standard curves, approximately 2ug was assayed per well of 96 well Agilent cell culture dishes (Agilent). The plate was run on an Agilent XFe96 Seahorse analyzer. Data was exported to Xcel and analyzed in GraphPad Prism.

### Echocardiography

Echocardiography was completed with a Vevo2100 ultrasound system at the University of Minnesota Imaging Center. M-mode images were used to measure tricuspid annular plane systolic excursion(7,10,19,36).

### Closed-chest pressure-volume hemodynamic studies

5% inhaled isoflurane was used for induction and then rats were maintained on 2-3% isoflurane while they were ventilated with a SomnoSuite small animal anesthesia system (Kent Scientific, Torrington, CT). A high-fidelity catheter (Scisense 1.9F pressure-volume, Transonic Systems, Ithaca, NY) was placed into the RV via the right internal jugular vein. RV pressure and volume were measured with the Transonic AV500 Pressure-Volume Measurement System and then analyzed on LabScribe version 4 (iWorx Systems, Dover, NH) as we have previously described(7,10,19).

### RV fibroblast isolation and culture

Fibroblasts were isolated and cultured according to Tian *et al(37)*. After two passages and expansion, 10 cm cell culture plates were rinsed with PBS and 400µl of lysis buffer with protease and phosphatase inhibitors was applied, collected and processed as described by Tian *et al*. Protein was quantified by BCA assay and Western blots were performed as described above. Loading control for these experiments was vimentin.

### Statistics

Statistical analyses were performed with GraphPad Prism 9.0 (San Diego, CA). Normality of data was determined by Shapiro-Wilk test. If data was normally distributed and there was equal variance as determined by the Brown-Forsythe test, one-way analysis of variance (ANOVA) with Tukey’s multiple comparisons test was performed. If there was unequal variance, Brown-Forsythe and Welch ANOVA with Dunnett multiple comparisons test was completed. If the data was not normally distributed, Kruskal-Wallis test and Dunn’s multiple comparisons test were used. Graphs show the mean value and all individual values.

Hierarchical cluster analyses and correlational heatmapping were performed with MetaboAnalyst software(38). Relative abundance of each protein in proteomics studies was determined with Scaffold (MCT experiments) or Proteome Discover Software (SuHx experiments) and those values were used for both analyses.

---

**Supplemental Table 1:** Antibodies and Dilutions Used in Study

| Antigen | Company | Catalogue Number | Dilution | Experimental Approach |
| --- | --- | --- | --- | --- |
| AMP kinase | Cell Signaling Technology | 2532 | 1:200 | Western blot |
| Phosphorylated AMP kinase | Cell Signaling Technology | 2531 | 1:200 | Western blot |
| α-tubulin | Millipore | 05-829 | 1:250 | Western blot |
| β-tubulin | Sigma-Aldrich | T4026 | 1:250 | Western blot |
| Detyrosinated α-tubulin | Abcam | ab48389 | 1:250 | Western blot |
| MAP4 | Millipore | ab6020 | 1:100 | Western blot |
| Phosphorylated MAP4 | Thermo Scientific | B5-5438R | 1:100 | Western blot |
| CLIP-170 | Cell Signaling Technology | 8977S | 1:250 | Western blot |
| Phosphorylated CLIP-170 | Kind gift from Dr. Yasunori Shintani | Not applicable | 1:250 | Western blot |
| Junctophilin-2 | Thermo Fisher Scientific | A304-455A | 1:250 | Western blot |
| Vimentin | R&D Systems | AF2105-SP | 1:250 | Western blot |
| β-tubulin | Abcam | Ab6046 | 1:50 | Immunofluorescence |
| Catalase | Abcam | Ab209211 | 1:50 | Immunofluorescence |
| Anti-rabbit Alexa 555 secondary antibody | Invitrogen | A32732 | 1:500 | Immunofluorescence |
| Anti-mouse secondary | Li-COR | 926-32210<br>926-68070 | 1:5000 | Western blot |
| Anti-rabbit secondary | Li-COR | 926-32211<br>926-68071 | 1:5000 | Western blot |

**Supplemental Figure 1:**
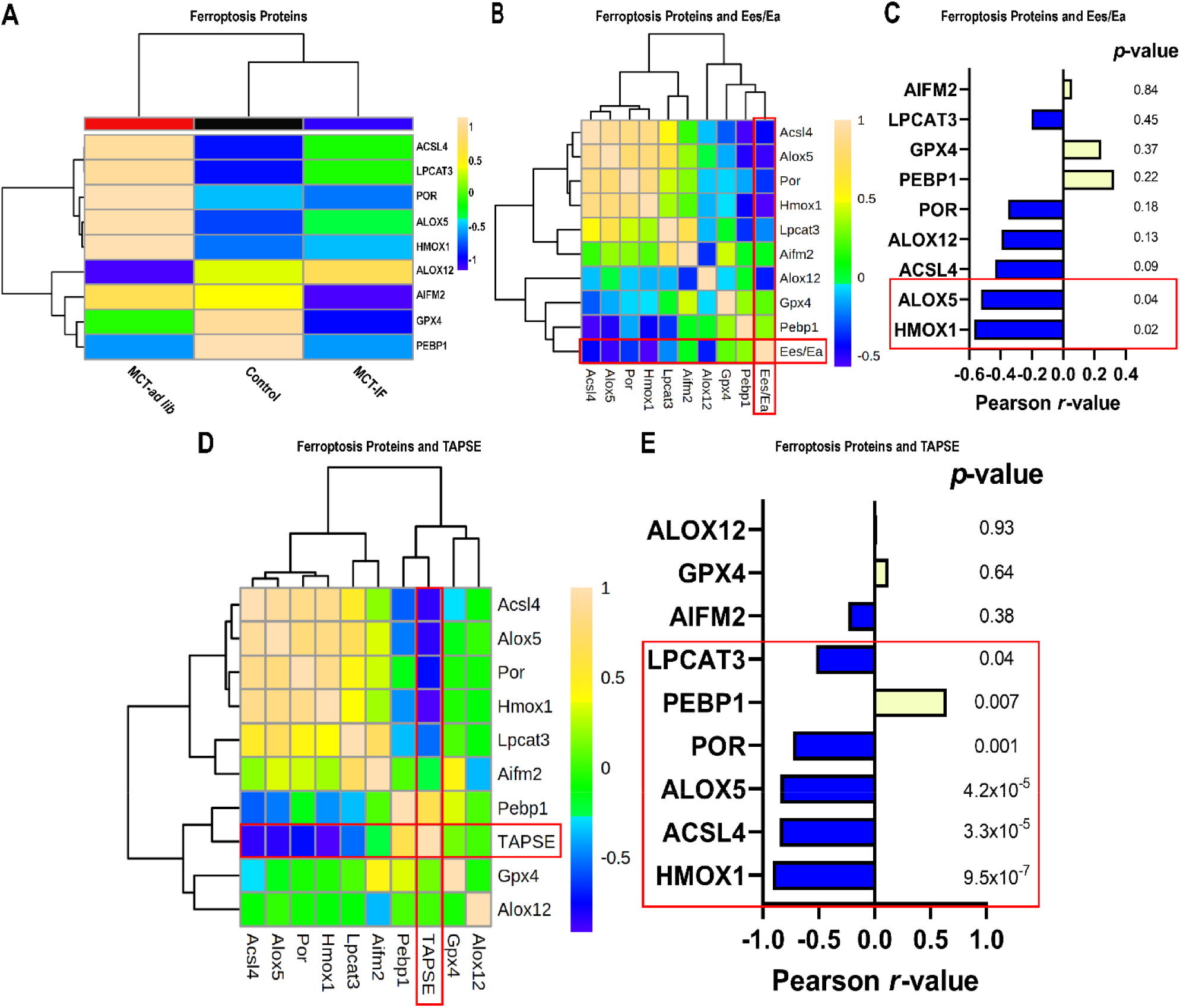
Ferroptosis proteins are dysregulated in RV failure and pro-ferroptotic proteins are associated with depressed RV function in MCT rats. (A) Hierarchical cluster analysis of ferroptosis proteins. MCT-*ad lib* RVs had higher levels of pro-ferroptotic proteins, which was mitigated by IF. (B) Correlational heat map of ferroptotic proteins and Ees/Ea. (C) *r*-value and *p*-value for each protein’s relationship with Ees/Ea. Red box highlights significant relationships. (D) Correlational heat map of ferroptotic proteins and TAPSE. (E) *r*-value and *p*-value for each protein’s relationship with TAPSE. Red box highlights significant relationships.

**Supplemental Figure 2:**
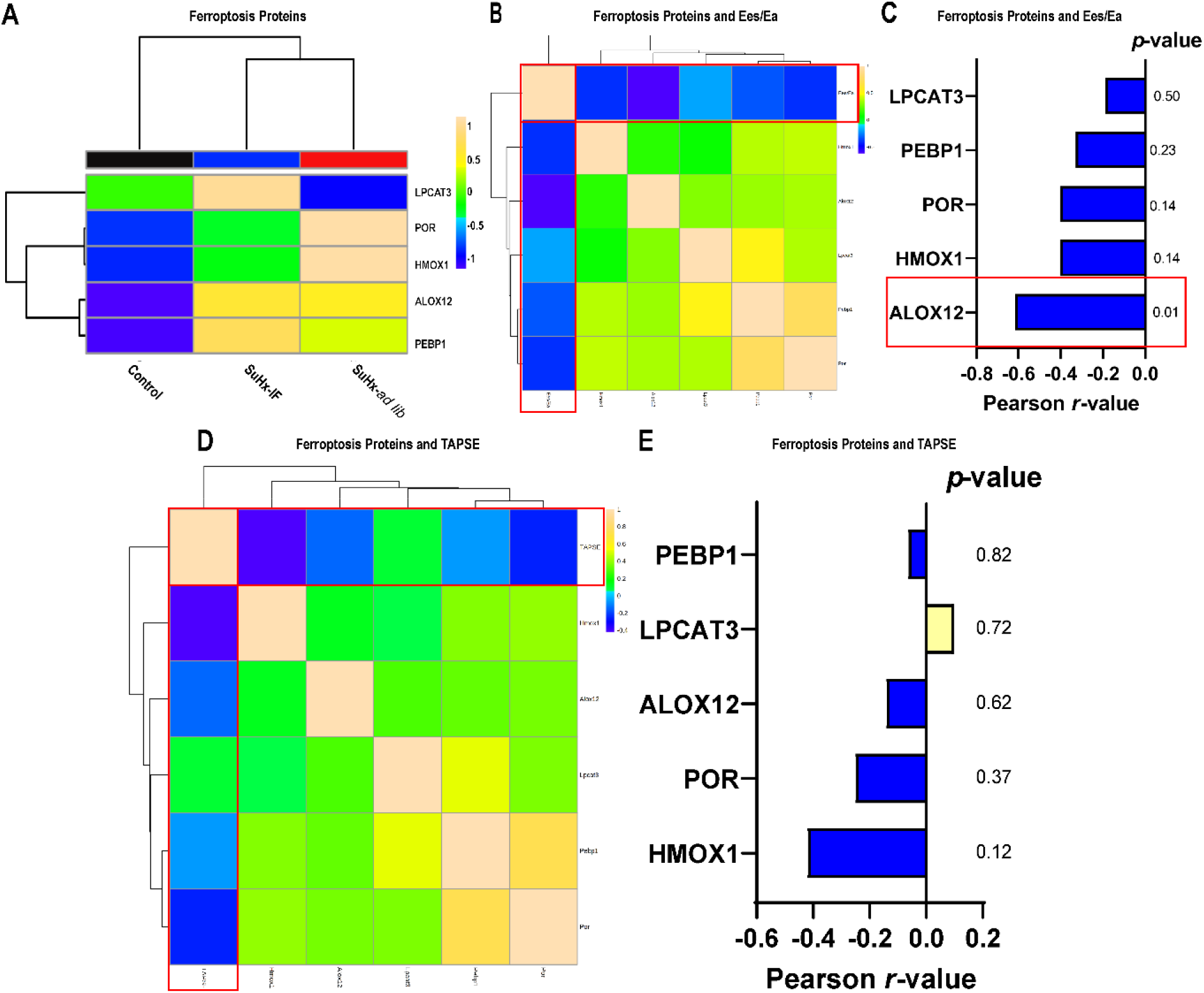
Ferroptosis proteins are upregulated in SuHx-mediated RV failure. (A) Hierarchical cluster analysis of ferroptosis proteins. SuHx-*ad lib* RVs had higher levels of pro-ferroptotic proteins, which was mitigated by IF. (B) Correlational heat map of ferroptotic proteins and Ees/Ea. (C) *r*-value and *p*-value for each protein’s relationship with Ees/Ea. Red box highlights significant relationships. (D) Correlational heat map of ferroptotic proteins and TAPSE. (E) *r*-value and *p*-value for each protein’s relationship with TAPSE. Red box highlights significant relationships.

**Supplemental Figure 3:**
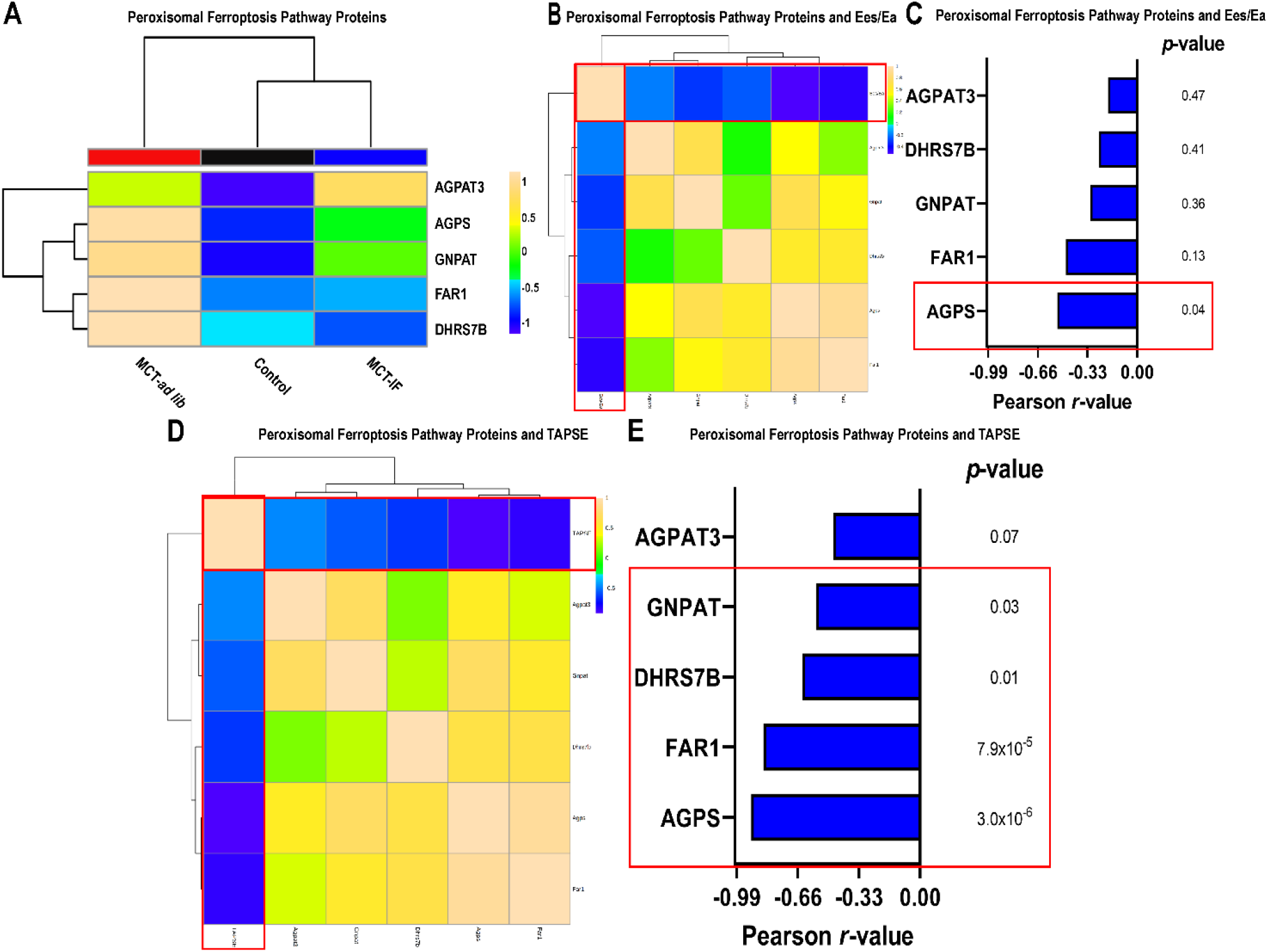
Peroxisomal-mediated ferroptosis proteins are upregulated in RV failure and inversely associated with RV function in MCT rats. (A) Hierarchical cluster analysis of ferroptosis proteins. MCT-*ad lib* RVs had higher levels of nearly all peroxisomal-mediated ferroptotic proteins, which was mitigated by IF. (B) Correlational heat map of peroxisomal-mediated ferroptotic proteins and Ees/Ea. (C) *r*-value and *p*-value for each protein’s relationship with Ees/Ea. Red box highlights significant relationships. (D) Correlational heat map of peroxisomal-mediated ferroptotic proteins and TAPSE. (E) *r*-value and *p*-value for each protein’s relationship with TAPSE. Red box highlights significant relationships.

**Supplemental Figure 4:**
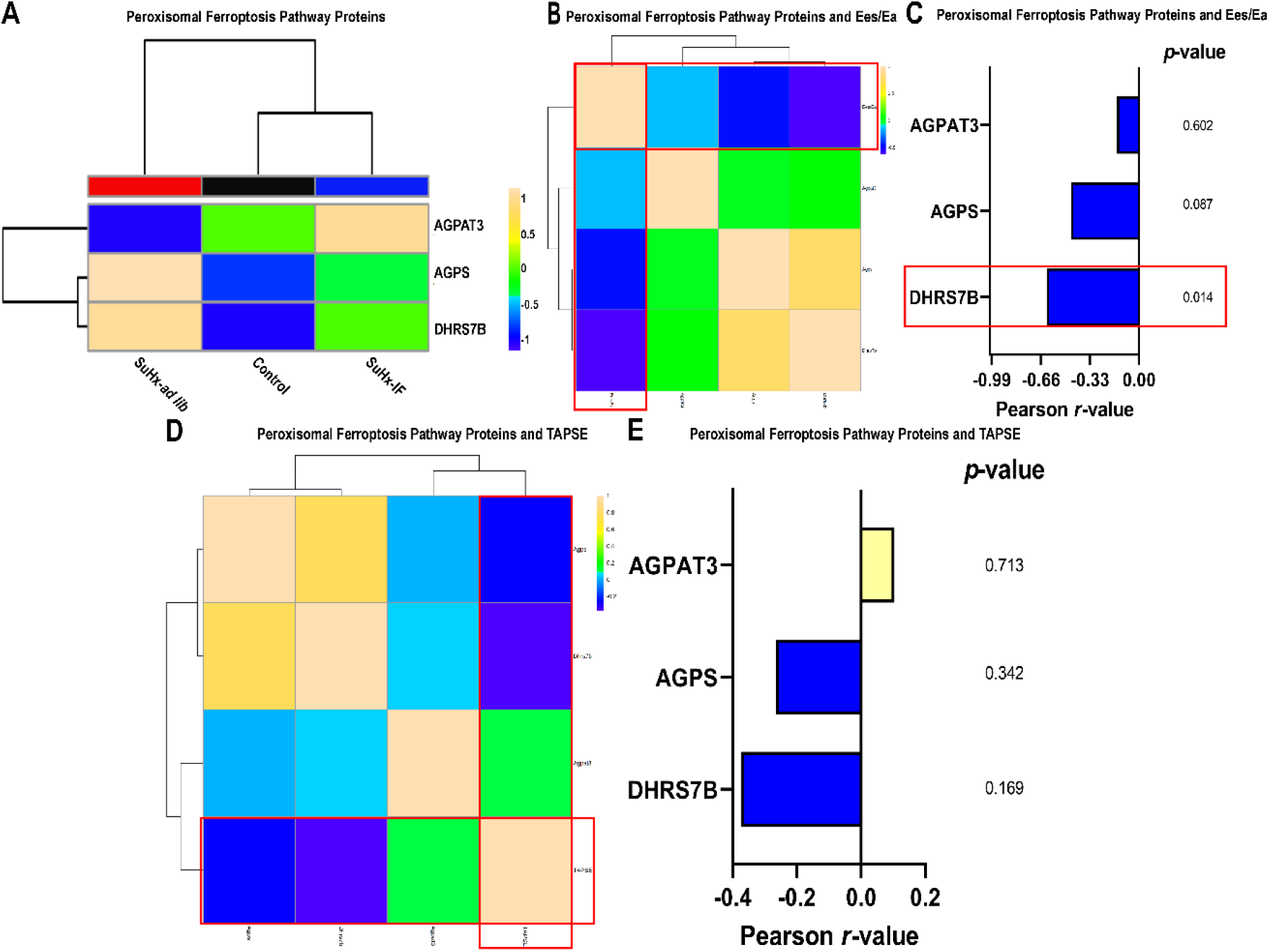
Some peroxisomal-mediated ferroptosis proteins are upregulated in Sugen-hypoxia rats, which was suppressed by IF. (A) Hierarchical cluster analysis of ferroptosis proteins. SuHx-*ad lib* RVs had higher levels of nearly all peroxisomal-mediated ferroptotic proteins, which was mitigated by IF. (B) Correlational heat map of peroxisomal-mediated ferroptotic proteins and Ees/Ea. (C) *r*-value and *p*-value for each protein’s relationship with Ees/Ea. Red box highlights significant relationships. (D) Correlational heat map of peroxisomal-mediated ferroptotic proteins and TAPSE. (E) *r*-value and *p*-value for each protein’s relationship with TAPSE. Red box highlights significant relationships.

**Supplemental Figure 5:**
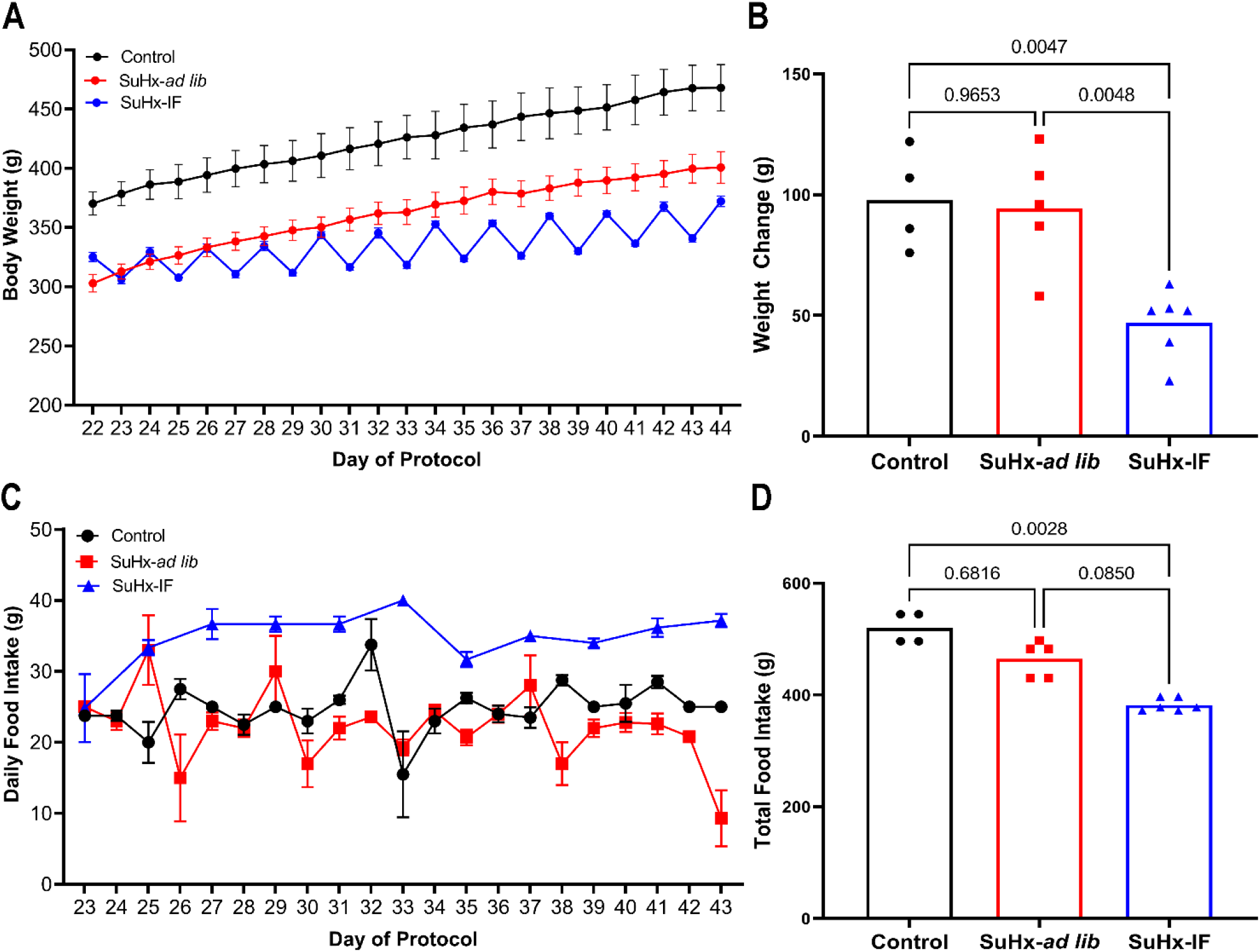
Analysis of food consumption and weight gain in Sugen-hypoxia experiments. (A) Change in body weight over time after three weeks of exposure to normoxia (controls) or hypoxia. (B) Total change in body weight over time under different experimental conditions. *p*-values as determined by one-way ANOVA with Tukey’s multiple comparisons test (C) Daily food consumption over duration of the experiment. (D) Total food consumption over the duration of the experiment. *p*-values as determined by Kruskal-Wallis test and Dunn’s multiple comparisons test.

**Supplemental Figure 6:**
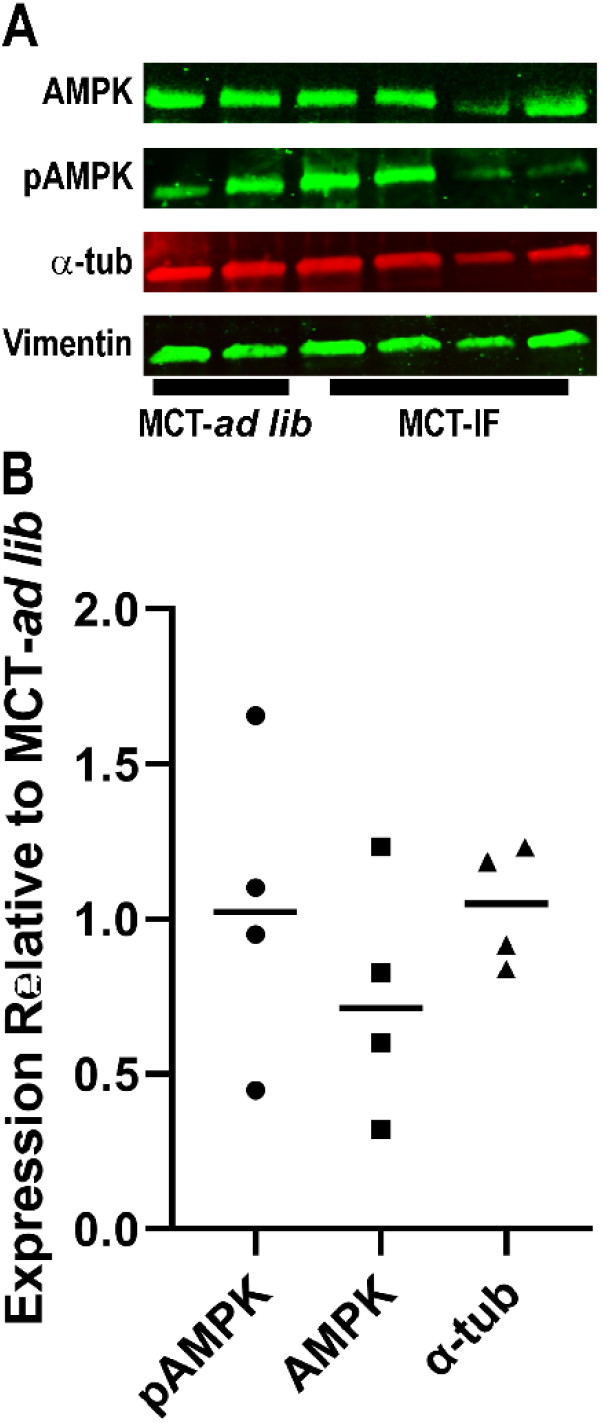
RV fibroblasts isolated from MCT rats did not exhibit differences in AMPK activation state or tubulin expression with intermittent fasting. (A) Representative Westerns blots and (B) quantification of protein levels from fibroblasts isolated from two MCT-*ad lib* and four MCT-IF RVs. Vimentin was used as the loading control.

**Supplemental Figure 7:**
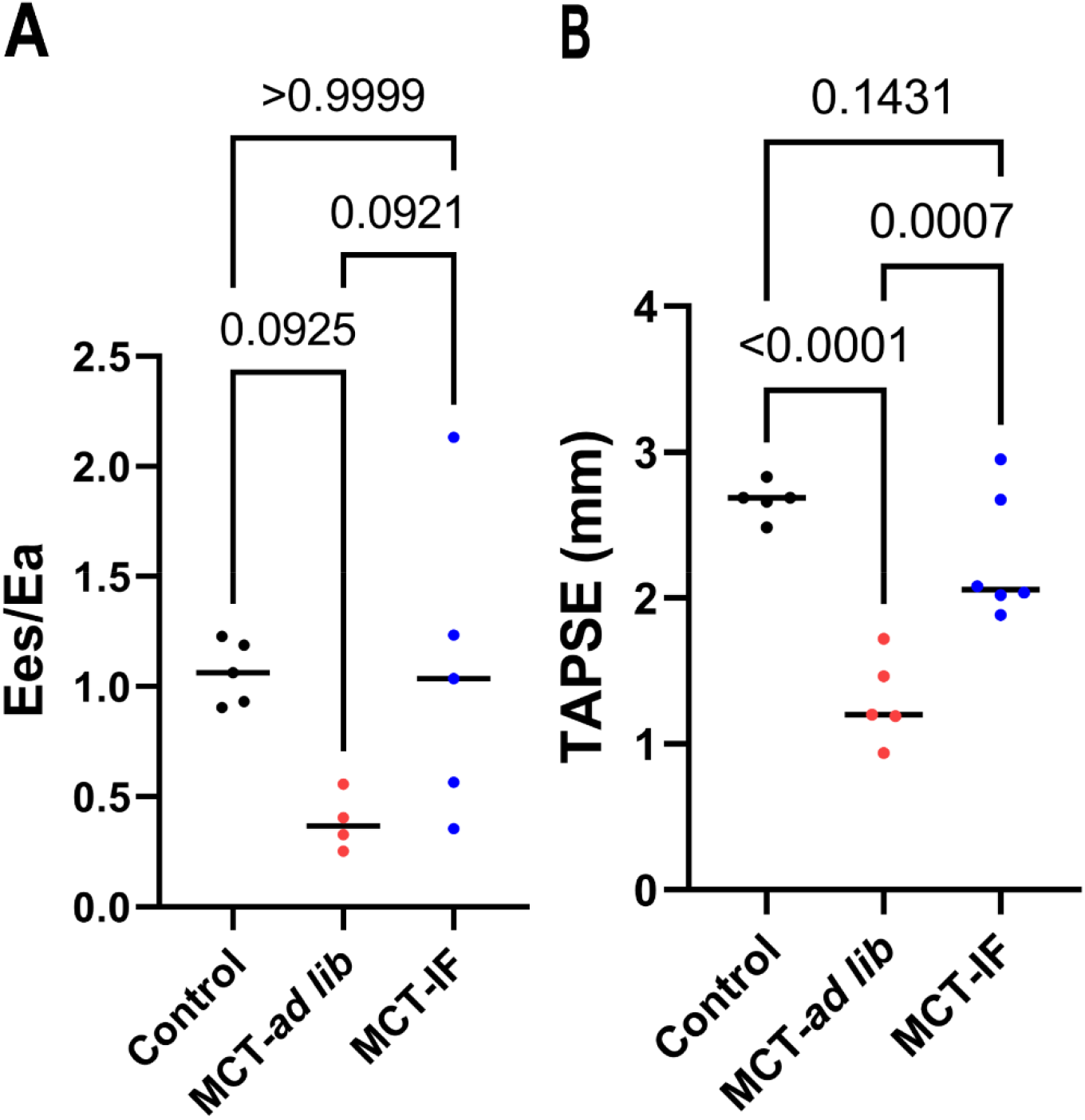
Quantification of RV function in MCT Experiments. Ees/Ea (A) and TAPSE (B) values obtained from the animals in MCT experiments that were analyzed in proteomics experiments. *p*-values as determined by one-way ANOVA with Tukey’s multiple comparisons test.

**Supplemental Figure 8:**
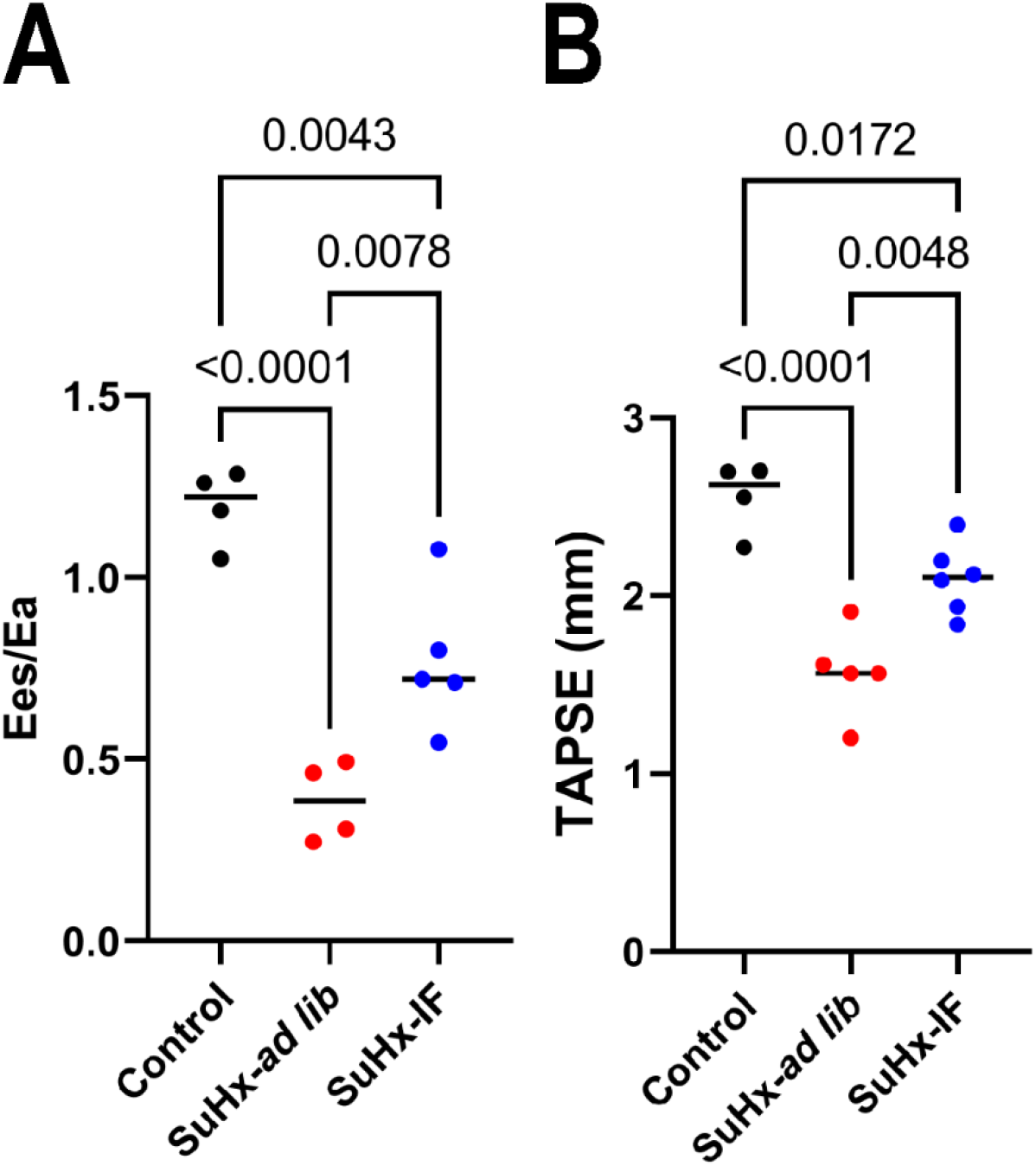
Quantification of RV function in Sugen-hypoxia experiments. (A) Ees/Ea and TAPSE (B) values from the animals evaluated in Sugen-hypoxia experiments. *p*-values as determined by one-way ANOVA with Tukey’s multiple comparisons test.

## Abbreviations

ACAA1: Peroxisome 3-ketoacyl-CoA thiolase
ACAD: Acyl-CoA dehydrogenase
ACSL: Acyl-CoA Synthetase Long Chain
AGPAT3: 1-acylglycerol-3-phosphate *O*-acyltransferase 3
AGPS: Alkylglycerone phosphate synthase
AMPK: Adenosine monophosphate-activated protein kinase
CLIP170: Cytoplasmic linker protein-170
Ees/Ea: End-systolic elastance/arterial elastance
EHHADH: L-bifunctional enzyme
ETC: Electron transport chain
FAR1: Fatty acyl-CoA reductase 1
GNPAT: Glyceronephosphate *O*-acyltransferase
IF: Intermittent fasting
MAP4: Microtubule-associated protein 4
MCT: Monocrotaline
PAH: Pulmonary arterial hypertension
PHYH: Phytanoyl-CoA 2-hydroxylase
pMAP4: Microtubule-associated protein 4 Ser941 phosphorylated
RKIP/PEBP1: Raf kinase inhibitor protein
RV: Right ventricular
SCP2: Sterol carrier protein 2
SuHx: Sugen-hypoxia
TAPSE: tricuspid plane annular systolic excursion
T-tubule: Transvere tubule

